# Plasticity of the Arabidopsis leaf lipidome and proteome in response to pathogen infection and heat stress

**DOI:** 10.1101/2024.02.09.579702

**Authors:** Patricia Scholz, Nathan M. Doner, Katharina Gutbrod, Cornelia Herrfurth, Philipp Niemeyer, Magdiel S. S. Lim, Katharina Blersch, Kerstin Schmitt, Oliver Valerius, John Shanklin, Ivo Feussner, Peter Dörmann, Gerhard H. Braus, Robert T. Mullen, Till Ischebeck

## Abstract

Plants must cope with a variety of stressors during their life cycle, and the adaptive responses to these environmental cues involve all cellular organelles. Among them, comparatively little is known about the contribution of cytosolic lipid droplets (LDs) and their core set of neutral lipids and associated surface proteins to the rewiring of cellular processes in response to stress. Here, we analyzed the changes that occur in the lipidome and proteome of Arabidopsis leaves after pathogen infection with *Botrytis cinerea* or *Pseudomonas syringae*, or after heat stress. Analyses were carried out in wild-type plants and the oil-rich double mutant *tgd1-1 sdp1-4* that allowed for an allied study of the LD proteome in stressed leaves. Using liquid chromatography-tandem mass spectrometry-based methods, we show that a hyperaccumulation of the primary LD core lipid triacylglycerol is a general response to stress and that acyl chain and sterol composition are remodeled during cellular adaptation. Likewise, comparative analysis of the LD protein composition in stress-treated leaves highlighted the plasticity of the LD proteome as part of the general stress response. We further identified at least two additional LD-associated proteins, whose localization to LDs in leaves was confirmed by confocal microscopy of fluorescent protein fusions. Taken together, these results highlight LDs as dynamic contributors to the cellular adaptation processes that underlie how plants respond to environmental stress.

**One sentence summary:** Biotic and heat stress strongly alters the lipidome and proteome of Arabidopsis leaves including the proteome of lipid droplets.

## Introduction

Plants naturally encounter a plethora of abiotic and biotic threats during their life cycle. Consequently, plant cells have to be highly adaptive at the transcriptomic, proteomic and metabolic level, which requires the interplay of different signaling pathways and organelles (Kumar et al., 2016; Zhu, 2016; Crawford et al., 2018). One pertinent example of this cellular interplay is lipid remodeling, whereby fatty acid synthesis and lipid turnover in the plastids, and lipid turnover in the endomembrane system act together to modify cellular membrane composition in response to stress. More specifically, acyl chains removed from the membrane lipid pool appear to be transferred into triacylglycerols (TAGs) stored in cytosolic lipid droplets (LDs; Xu and Shanklin, 2016). In heat-stressed Arabidopsis seedlings for example, cytosolic TAGs with a high degree of unsaturation accumulate and the respective polyunsaturated acyl chains originate, at least in part, from chloroplasts (Mueller et al., 2015; Mueller et al., 2017). Similarly, in tobacco pollen tubes, heat stress leads to an increase in the proportion of saturated acyl chains in the membrane lipids phosphatidylcholine (PC) and phosphatidylethanolamine (PE), at the expense of mono- and polyunsaturated acyl chains, while total TAG levels increase (Krawczyk et al., 2022a). Similar effects have been observed in heat-stressed leaves of Arabidopsis (Higashi et al., 2015) and accumulation of TAG is linked to a number of stresses, e.g., freezing stress (Moellering et al., 2010), drought and cold treatment (Tarazona et al., 2015) or pathogen infection (Schieferle et al., 2021), which implies that LDs are important organelles in the plant stress response.

Cytosolic LDs consist of a hydrophobic core of neutral lipids, primarily TAGs and sterol esters (SEs), delimited by a monolayer of phospholipids (Guzha et al., 2023). Embedded into and/or associated with the surface of the monolayer are various proteins that convey to the LDs different functions depending on the cellular context (Brocard et al., 2017; Ischebeck et al., 2020; Kretzschmar et al., 2020). Most studies on LDs have been carried out with tissues/organs where they are highly abundant, including seeds, seedlings and pollen (Vance and Huang, 1987; Tzen et al., 1993; Chen et al., 1999; Lin et al., 2002; Hsieh and Huang, 2004; Kretzschmar et al., 2020). Indeed, the best described LD proteomes are those in oilseeds, where members of the oleosin (OLE), caleosin (CLO) and steroleosin (also called hydroxysteroid dehydrogenase [HSD]) protein families predominate (Jolivet et al., 2004; Katavic et al., 2006; Jolivet et al., 2009; Kretzschmar et al., 2018; Kretzschmar et al., 2020).

LDs function in seeds predominantly as lipid storage organelles, and oleosins are considered to function primarily in stabilizing LDs and preventing their coalescence (Cummins et al., 1993; Murphy, 1993). However, during seedling establishment, the LD proteome undergoes a transformation as the seed LD proteins are degraded and other LD proteins confer new functionalities to the LDs (Deruyffelaere et al., 2018; Kretzschmar et al., 2018; Kretzschmar et al., 2020). Among the latter, the LD-ASSOCIATED PROTEIN (LDAP) family, the LDAP-INTERACTING PROTEIN (LDIP), and SEIPIN proteins, which are endoplasmic reticulum (ER) membrane proteins that are situated at ER-LD junctions, are crucial for LD biogenesis (Cai et al., 2015; Taurino et al., 2018; Pyc et al., 2021). Additional LD proteins that function in seedlings have also been described, including those with enzyme activities involved in the synthesis or breakdown of primary and secondary metabolites (Corey et al., 1993; Diener et al., 2000; Shimada et al., 2014a; Müller and Ischebeck, 2018). Furthermore, a protein partially localizing to LDs, RABC1/LDS1, influences LD dynamics during guard cell development (Ge et al., 2022).

Environmental changes also influence LD-related processes. LD abundance in Arabidopsis leaves, for instance, increases in response to drought, cold or heat stress (Gidda et al., 2016; Doner et al., 2021). Correspondingly, Arabidopsis plants that over-accumulate LDs, such as transgenic lines overexpressing *LDAPs* (Gidda et al., 2016), are also more drought-tolerant (Kim et al., 2016a), further suggesting that LDs play a role in stress tolerance. In addition, CLO3 is probably the best characterized LD protein in non-seed tissues and has been implicated in stress responses in leaves (Partridge and Murphy, 2009; Aubert et al., 2010; Blée et al., 2014). For example, a role of CLO3 in the biotic stress response has been attributed to its peroxygenase function (Shimada et al., 2014a). After infection of Arabidopsis leaves with the fungal pathogen *Colletotrichum higginsianum*, CLO3 and α-DIOXYGENASE 1 (α-DOX1) accumulate at LDs in the perilesional area of the infection. There, they are thought to serve in tandem to convert α-linolenic acid, stored in the neutral lipids of LDs, into 2-hydroxy-octadecatrienoic acid (2-HOT), which then counteracts fungal spread (Shimada et al., 2014a).

Despite the growing knowledge on selected LDs proteins and their roles in vegetative plant organs, the relatively low abundance of LDs in these tissues has generally been a technical challenge for lipidomic and/or proteomic studies. Nevertheless, successful studies have been carried out on the leaf LD proteome of senescing and drought-stress leaves, as well as leaves infected with *Pseudomonas syringae* pv. *tomato* (*Pto*) DC3000 *avrRpm1* (Brocard et al., 2017; Fernández-Santos et al., 2020; Doner et al., 2021), which have resulted in the identification of several novel LD proteins. However, the dynamics of LDs, especially in terms of their proteome, during the plant stress response remain unclear. Leaves are constantly exposed to a vast array of environmental conditions, so it seems likely that the composition of leaf LDs is highly flexible in order to react to these external cues. Here, we assessed the changes that leaves and leaf LDs undergo when subjected to biotic and abiotic stresses. Arabidopsis plants were infected with one of two different pathogens or exposed to heat stress, and subsequently their leaf proteome and lipidome were analyzed. Comparisons with control treatments allowed us to observe the alterations induced by the three different treatments. In addition, proteomic analysis of LD-enriched fractions isolated from leaves subjected to the same treatments enabled us to survey specifically the dynamics of the LD proteome and, in doing so, identify two new LD proteins.

## Results

### Heat stress and pathogen infection cause differential changes in the Arabidopsis leaf lipidome

Reports on TAG accumulation and elevated numbers of LDs induced by abiotic and biotic stresses were among the first indications for the involvement of LDs in the stress response of leaves (Gidda et al., 2016; Higashi and Saito, 2019; Doner et al., 2021; Schieferle et al., 2021). To confirm these previous observations and determine if the overall lipidome is also affected by our treatments, we analyzed changes in the leaf lipidome in infected or heat-stressed plants (Suppl. Datasets S1 - S7). For this, we subjected seven-week-old Arabidopsis wild-type plants to heat stress (consisting of 24 hours at 37°C) or spray-infected them with either *Botrytis cinerea* or *Pseudomonas syringae* pv. *tomato* DC3000 *ΔavrPto/ΔavrPtoB* (hereafter: *Pseudomonas*), and subsequently carried out lipidomic analyses.

Overall, most of the analyzed lipid classes did not change significantly in abundance. However, triacylglycerol (TAG) levels were significantly higher after *Pseudomonas* infection and heat treatment and were also increased, albeit not significantly, after infection with *B. cinerea* (Figure 1). Interestingly, the amount of sterol esters (SEs) decreased under heat stress. Furthermore, several lipid classes showed an altered distribution of molecular lipid species (Figures 2 and 3; Suppl. Figures S1 – S4), most prominently for heat stressed leaves with increased proportions of acyl chains with a lower number of carbon atoms and fewer double bonds in membrane lipids like mono- and digalactosyldiacylglycerol (MGDG, DGDG, Figure 2) and phosphatidylcholine and phosphatidylethanolamine (PC, PE). In contrast, in TAGs of heat-stressed leaves, lipid species with a higher number of carbon atoms and more double bonds increased in proportion. For infection treatments, acyl chain adaptations likewise occurred albeit with a less pronounced pattern (Figure 3). Interestingly, both *Pseudomonas* and *B. cinerea* infections led to similar changes in the profile of free sterols, as the contribution of stigmasterol and isofucosterol increased at the expense of β-sitosterol (Figure 3), while the effects of heat stress on free sterols were less prominent (Suppl. Figure S1).

**Figure 1:**
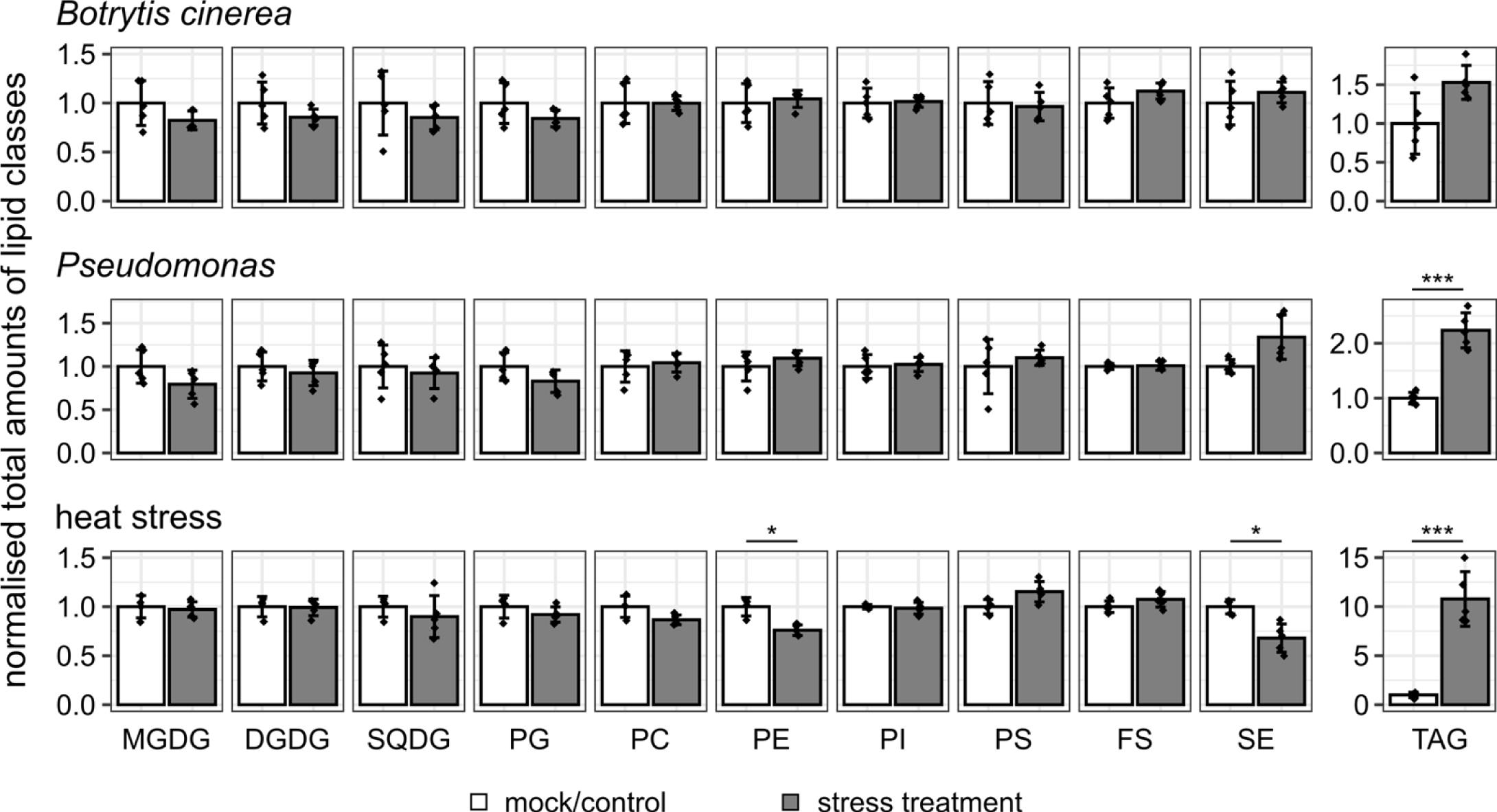
Changes in total abundance of membrane and hydrophobic lipids after different stress treatments. Arabidopsis plants were infected with *Botrytis cinerea* or *Pseudomonas syringae* pv. *tomato* DC3000 *ΔavrPto/ΔavrPtoB* (*Pseudomonas*), or kept for 24 h at 37 °C (heat stress) and compared to mock-infected or non-stressed plants. After stress treatment, leaves were harvested, lipids isolated, and their amounts determined by mass spectrometry. Values of all species in the indicated lipid classes were added up and this sum was normalized to the respective control. Statistical comparisons were calculated with Student’s t-test, using Holm-Bonferroni correction for multiple comparisons. Values are shown as mean ± standard deviation. Significant differences are indicated with * and *** for p<0.05 and p<0.001, respectively. n ≥ 4 biological replicates. MGDG, monogalactosyldiacylglycerol; DGDG, digalactosyldiacylglycerol; SQDG, sulfoquinovosyldiacylglycerol; PG, phosphatidylglycerol; PC, phosphatidylcholine, PE, phosphatidylethanolamine; PI, phosphatidylinositol; PS, phosphatidylserine; FS, free sterols; SE, sterol esters; TAG, triacylglycerol.

**Figure 2:**
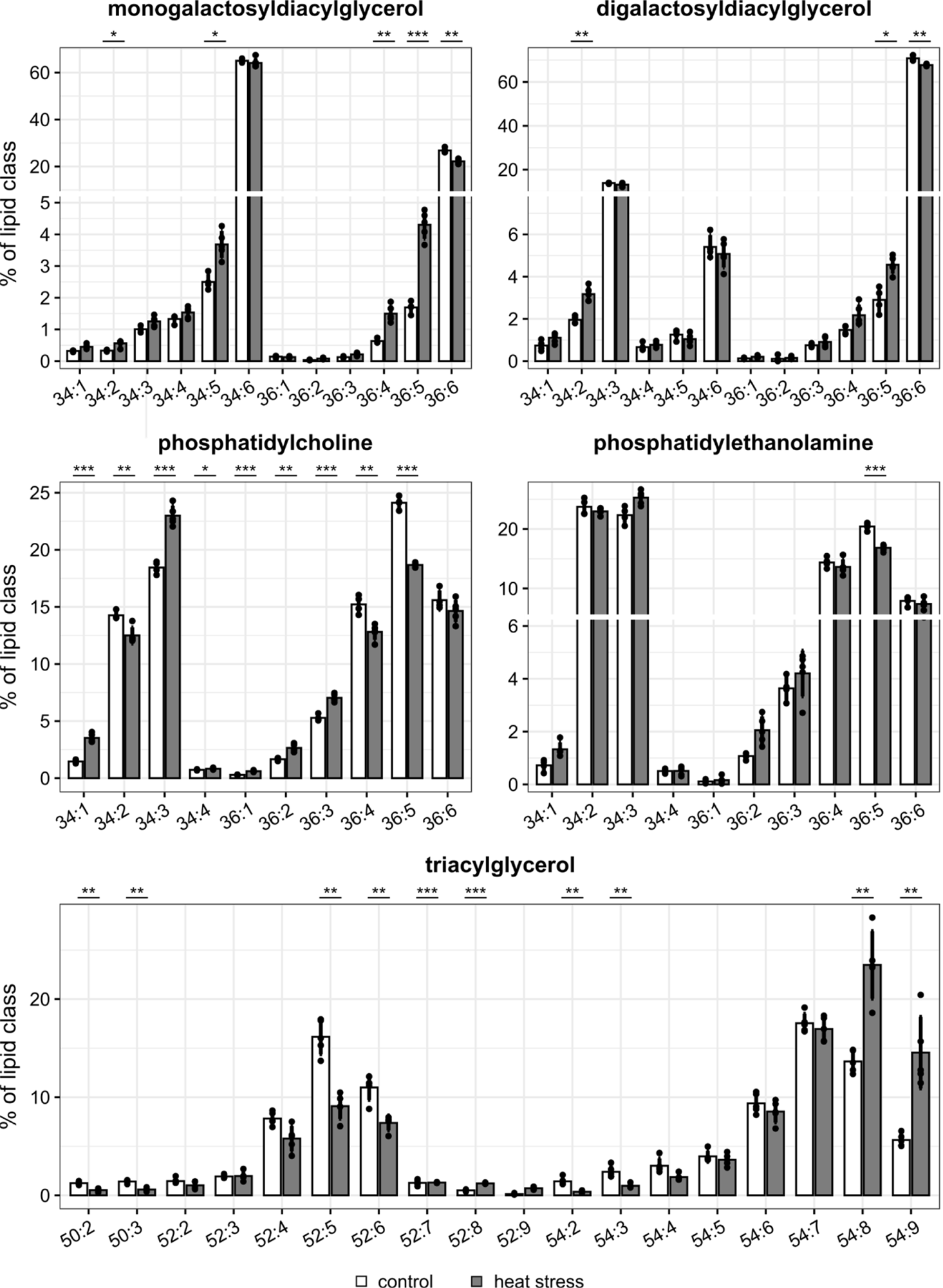
Heat-induced changes in membrane lipid and triacylglycerol composition. Arabidopsis Col-0 was heat-stressed (37°C for 24 h) or kept at normal temperature regime (control). Subsequently, membrane and hydrophobic lipids were isolated and analyzed by mass spectrometry. For individual lipids classes, the relative species composition was calculated and is shown here for the glycoglycerolipids monogalactosyldiacylglycerol and digalactosyldiacylglycerol, the phosphoglycerolipids phosphatidylcholine and phospatidylethanolamine, and triacylglycerol. Lipid species are described by the combined number of all carbon atoms and double bonds of all fatty acids esterified to the glycerol backbone. Statistical comparisons were calculated with Student’s t-test, using Holm-Bonferroni correction for multiple comparisons. Values are shown as mean ± standard deviation. Significant differences are indicated with *, ** and *** for p < 0.05, p < 0.01 and p < 0.001, respectively. n ≥ 4 biological replicates.

**Figure 3:**
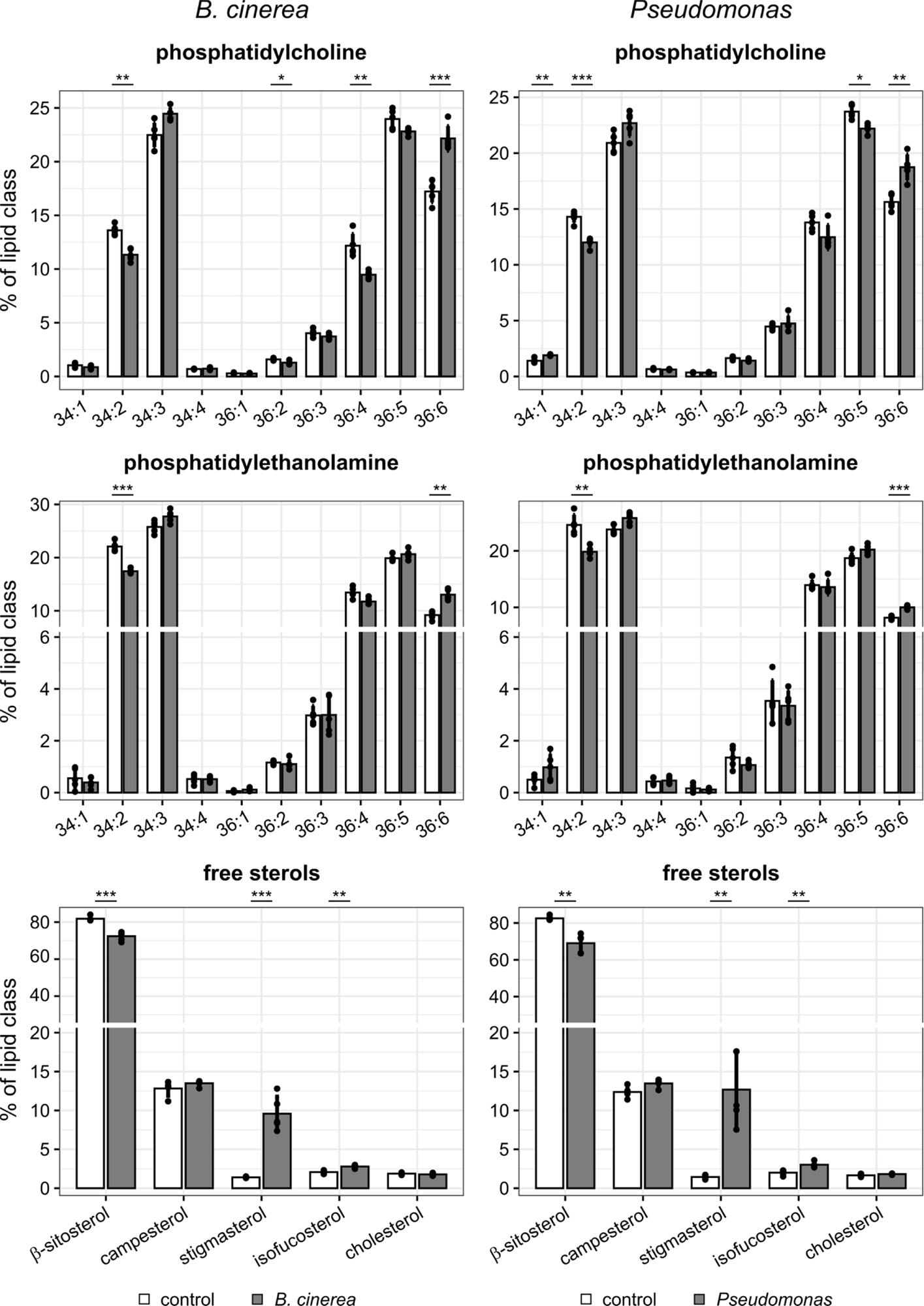
Alterations in the Arabidopsis lipid profile of phosphatidylcholine, phosphatidylethanolamine and free steroles after infection. Arabidopsis Col-0 plants were infected with *Botrytis cinerea* or *Pseudomonas syringae* pv. *tomato* DC3000 *ΔavrPto/ΔavrPtoB* (*Pseudomonas*). After the infection, lipids were isolated from leaves and analyzed by mass spectrometry. The relative composition of lipid species was determined and is displayed here for phosphatidylcholine, phosphatidylethanolamine and free sterols. For the phosphoglycerolipids, lipid species are described by the combined number of all carbon atoms and double bonds of all fatty acids esterified to the glycerol backbone. Statistical comparisons were calculated with Student’s t-test, using Holm-Bonferroni correction for multiple comparisons. Values are shown as mean ± standard deviation. Significant differences are indicated with *, ** and *** for p < 0.05, p < 0.01 and p < 0.001, respectively. n ≥ 4 biological replicates.

### The Arabidopsis leaf proteome changes in response to biotic and heat stress

To test if the proteome also undergoes changes, we extracted the total proteome from Arabidopsis leaves subject to the same stress treatments. Proteins from the mutant line *tgd1-1 sdp1-4,* that was stressed in parallel, were also extracted. Additionally, LD-enriched fractions were isolated. We used the *tgd1-1 sdp1-4* double mutant, as it has increased TAG levels stored in LDs in leaves (Fan et al., 2014) thereby making it easier to isolate LDs especially under unstressed conditions. At the same time, TAG accumulation in this mutant is not triggered by transcription factors that have a direct influence on gene expression like WRINKLED1 or LEAFY COTYLEDON 2 (Cernac and Benning, 2004; Kim et al., 2015; Qiao et al., 2022). For all samples, peptides were analyzed by liquid chromatography-tandem mass spectrometry (LC-MS/MS) and MS raw data was processed with the MaxQuant software (Cox and Mann, 2008) to identify and relatively quantify the original proteins. Two algorithms were used for quantification: the intensity-based absolute quantification (iBAQ) and the label-free quantification (LFQ) algorithm (Cox and Mann, 2008; Schwanhäusser et al., 2011; Cox et al., 2014). Protein quantification values were then normalized as per thousand of the total combined intensity in each sample, resulting in relative iBAQ (riBAQ) and relative LFQ (rLFQ) values (Suppl. Dataset S8, all metadata can be found in Suppl. Table S1). rLFQ values were used for comparisons of samples with similar protein compositions, i.e., total extract (TE) samples of different conditions or genotypes. riBAQ values were used for calculations of enrichment factors between LD-enriched fractions and TE fractions, as here sample composition differed strongly. Overall, this data set allowed us to compare (i) the effects of the different stresses on the total proteome in the wild type and double mutant line, (ii) differences between the wild type and the *tgd1-1 sdp1-4* double mutant, (iii) protein abundances in total cellular fractions and LD fractions to identify previously unknown LD-associated proteins (iv) changes especially in the LD proteome under stress in the double mutant.

Based on rLFQ values we compared changes of protein abundance in Col-0 leaves and calculated statistical significance of changes between the different treatments and their respective controls (Suppl. Datasets S9 – S11). The results were visualized as volcano plots, and proteins whose abundance was changed at least 1.5-fold between conditions with *p* < 0.05 were selected for further analysis (Figure 4).

**Figure 4:**
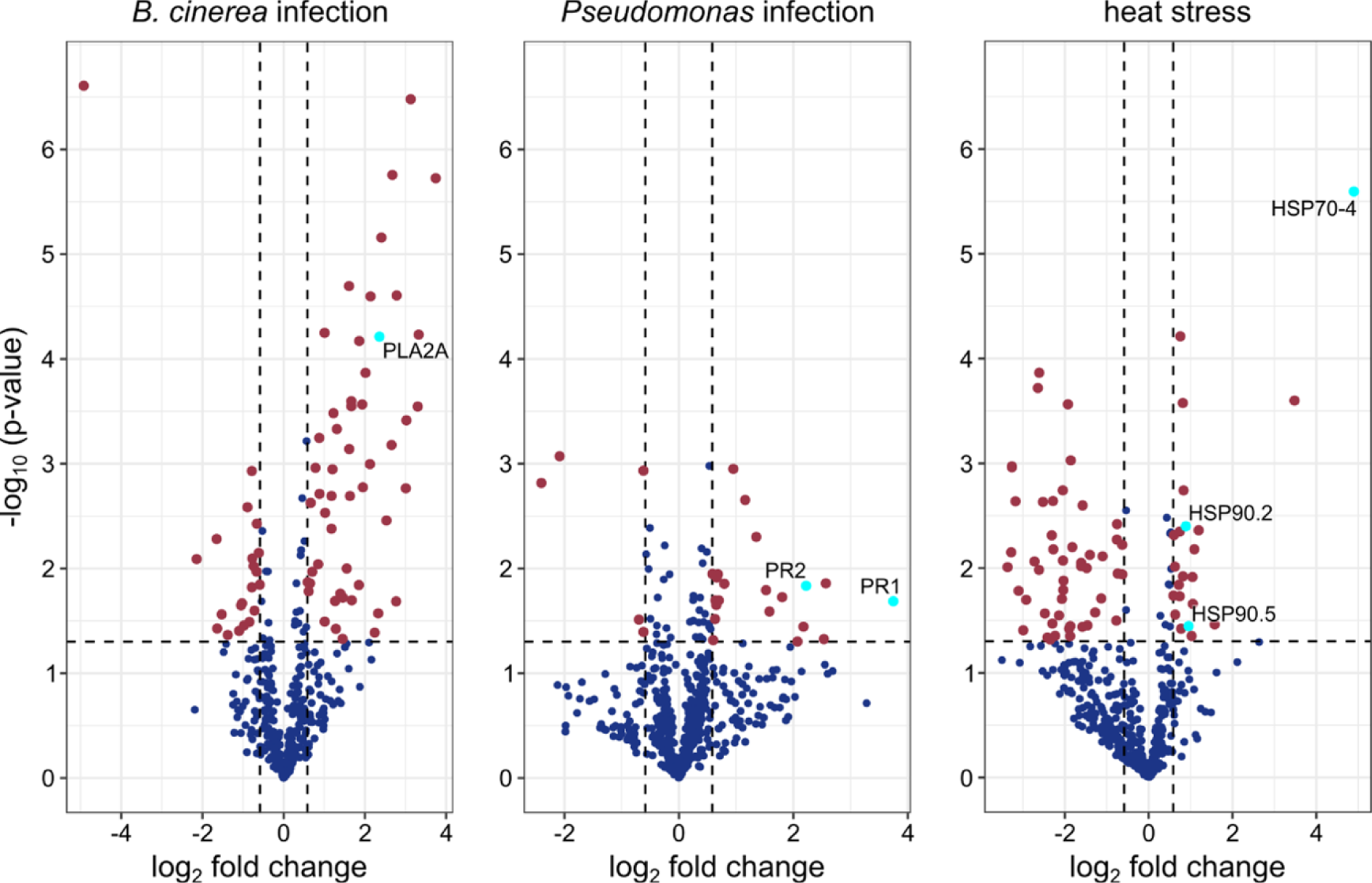
Alterations of total cellular proteins of Arabidopsis leaves subjected to different stress treatments. Arabidopsis plants were infected with *Botrytis cinerea* (*B. cinerea*), *Pseudomonas syringae* pv. *tomato* DC3000 *ΔavrPto/ΔavrPtoB* (*Pseudomonas*) or heat stressed for 24 h at 37 °C. Protein abundances (rLFQ values) of individual proteins were normalised to the respective values of the control treatment and the resulting ratio was log2-transformed. Statistical significance of the log2-fold change was calculated by Student’s t-test. The constructed volcano plots indicate proteins that are significantly enriched (upper right) or depleted (upper left) in reaction to the individual stress treatments. For each experiment, only proteins detected in all replicates of either mock-treated or infected plants were included in the analysis. For heat stress, proteins present in at least four replicates of either heat-stressed or control plants were analyzed. Vertical lines indicate 1.5-fold enrichment or depletion, while the horizontal line indicates a significance of *p* = 0.05. Proteins further mentioned in the text are labeled and highlighted in cyan. n=5 (biological replicates) for *B. cinerea* and its mock control, n=3 for Pseudomonas treatment and its control, n=5 for heat stress treatment and n=4 for its control.

Among the selected proteins, we identified individual proteins that had been linked previously to the respective stress treatment. For instance, for *B. cinerea* infection, the phospholipase PLA2A was only detected in our proteomic dataset after infection (Figure 4; Suppl. Dataset S9), which is consistent with a previous study that showed its accumulation after infection and dependent on jasmonic acid (JA) signaling (La Camera et al., 2005). Among the most highly increased proteins after *Pseudomonas* infection were the two pathogenesis-related (PR) proteins PR1 and PR2 (Suppl. Dataset S10), both of which are known to be upregulated as part of systemic acquired resistance to pathogen infection (Uknes et al., 1993; Fu and Dong, 2013). A third PR-protein, PR5, also more than doubled in abundance. Another defence-related protein that increased upon *Pseudomonas* infection was HYPERSENSITIVE INDUCED REACTION 2 (HIR2; Suppl. Dataset S7), which organizes immune receptors at the plasma membrane into nanoclusters (Qi et al., 2011; Qi and Katagiri, 2012). Finally, in response to heat treatment, the heat shock protein HSP70-4 showed the most pronounced increase in protein abundance (Figure 4), consistent with its role in thermotolerance against long term heat stress (Wang et al., 2021). Other heat shock proteins like HSP90.2 and HSP90.5 increased in abundance (Figure 4, Suppl. Dataset S11) as well, confirming that the plants were able to sense and respond to the applied temperature conditions.

In order to determine if whole networks of interacting and/or functionally-related proteins are changed under stress, we employed the web tool at the STRING v11.5 database (https://string-db.org; (Szklarczyk et al., 2021) to assess the differentially abundant proteins for each treatment with at least a 1.5-fold change and *p* < 0.05. Proteins with decreased or increased abundance were evaluated separately and only interactions of high confidence were analyzed. In plants stressed with *B. cinerea* or heat, chloroplastic and photosynthetic proteins dominated amongst proteins with decreased abundance (Suppl. Figure S5). Among the proteins with decreased abundance after *B. cinerea* infection were different members of photosystem I, II and the light-harvesting complexes. Similarly, after heat treatment, proteins of photosystem II were decreased as well as proteins of the chloroplastic electron transport chain and subunits of RuBisCO.

The upregulated proteins of the different treatments did not form as extensive networks as the downregulated ones (Suppl. Figure S5). Among the most extensive interaction networks upon heat stress was the one formed by heat shock proteins and chaperone proteins with the functional role of assisting protein folding. Other protein networks mostly contained two or three proteins, nevertheless, these small networks pointed towards metabolic adjustments induced by the different stress treatments. For example, heat treatment induced the accumulation of two catalases, CAT2 and CAT3, (Shimada et al., 2014a; Gidda et al., 2016) while infection with *B. cinerea* or *Pseudomonas* both led to an accumulation of glutathione S-transferase proteins. More specific for *B. cinerea*, metabolic adaptation included the upregulation of proteins involved in the biosynthesis of tryptophan and the detoxification of cyanide as for example TRYPTOPHAN SYNTHASE ALPHA CHAIN, TRYPTOPHAN SYNTHASE BETA-SUBUNIT 1, ß-CYANOALANINE SYNTHASE C1 and NITRILASE 4 (Yamaguchi et al., 2000; Piotrowski et al., 2001).

Among the 117 proteins with a known or putative function in lipid metabolism (Suppl. Dataset S12), only few were significantly changed, such as peroxisomal 3-ketoacyl-CoA thiolase, which is involved in β-oxidation (Germain et al., 2001) and was 2-fold upregulated under heat stress.

### The Arabidopsis *tgd1-1 sdp1-4* double mutant is altered in the leaf lipidome and proteome

As mentioned, LDs have been implicated with both biotic and abiotic stress responses (Shimada et al., 2014a; Gidda et al., 2016; Kim et al., 2016b; Fernández-Santos et al., 2020), and we also found strong increases of TAG under stress in our experiments. Given this, we investigated the LD proteome of Arabidopsis leaves under stress making use of the *tgd1-1 sdp1-4* double mutant. TAG levels in this mutant were reported to reach approximately 8% of leaf dry weight with a concomitant increase in number and size of leaf LDs (Fan et al., 2014). We could confirm a 65-fold increase of TAG levels in leaves of the *tgd1-1 sdp1-4* double mutant (Suppl. Figures S6). The acyl chain composition of leaf TAG in the mutant was also strongly altered, favoring TAGs with 54 carbon atoms, whose proportion increased from ca. 50% to ca. 70%. Interestingly though, the relative proportion of the most desaturated TAG species 54:9 decreased (Suppl. Figure S6). For SEs, a similar decrease of SE species with the acyl chain 18:3 could be observed, and the overall amounts of SEs decreased by ca. 25 % (Suppl. Figures S6, S7). Among membrane lipids, the mutant leaves contained significantly less of the plastidial lipids MGDG and DGDG while PC and phosphatidylinositol (PI) were significantly increased. Further, the acyl chain composition followed the trend already observed for TAGs and SEs, with generally decreased percentages of more highly desaturated lipid species (Suppl. Figure S6, S7). For example, in MGDG, DGDG, PC and PE, the proportions of 36:6 species were strongly decreased. In contrast, the phytosterol composition was not affected (Suppl. Figure S7).

When we examined the changes in the proteome of *tgd1-1 sdp1-4* mutant plant leaves under stress, we observed some common trends with the wild type (Suppl. Datasets S13-S15). This included for example a decrease in plastidial and photosynthetic proteins in reaction to stress and also the treatment-dependent accumulation of individual proteins described in the previous section (Suppl. Datasets S9-S11). However, when stress-responsive proteins were selected with the same criteria as for the wild type, less than 50% of proteins were shared amongst the up- or downregulated proteins of each treatment. As these differences pointed to underlying changes in the proteome of the double mutant, we decided to compare the proteome of *tgd1-1 sdp1-4* to the wild type under non-stressed conditions (Figure 5A, Suppl. Dataset S16). Using the same selection criteria as for the analysis of treatment-induced changes (enrichment or depletion of at least 1.5-fold, *p* < 0.05), 253 affected proteins were selected for further analysis.

**Figure 5:**
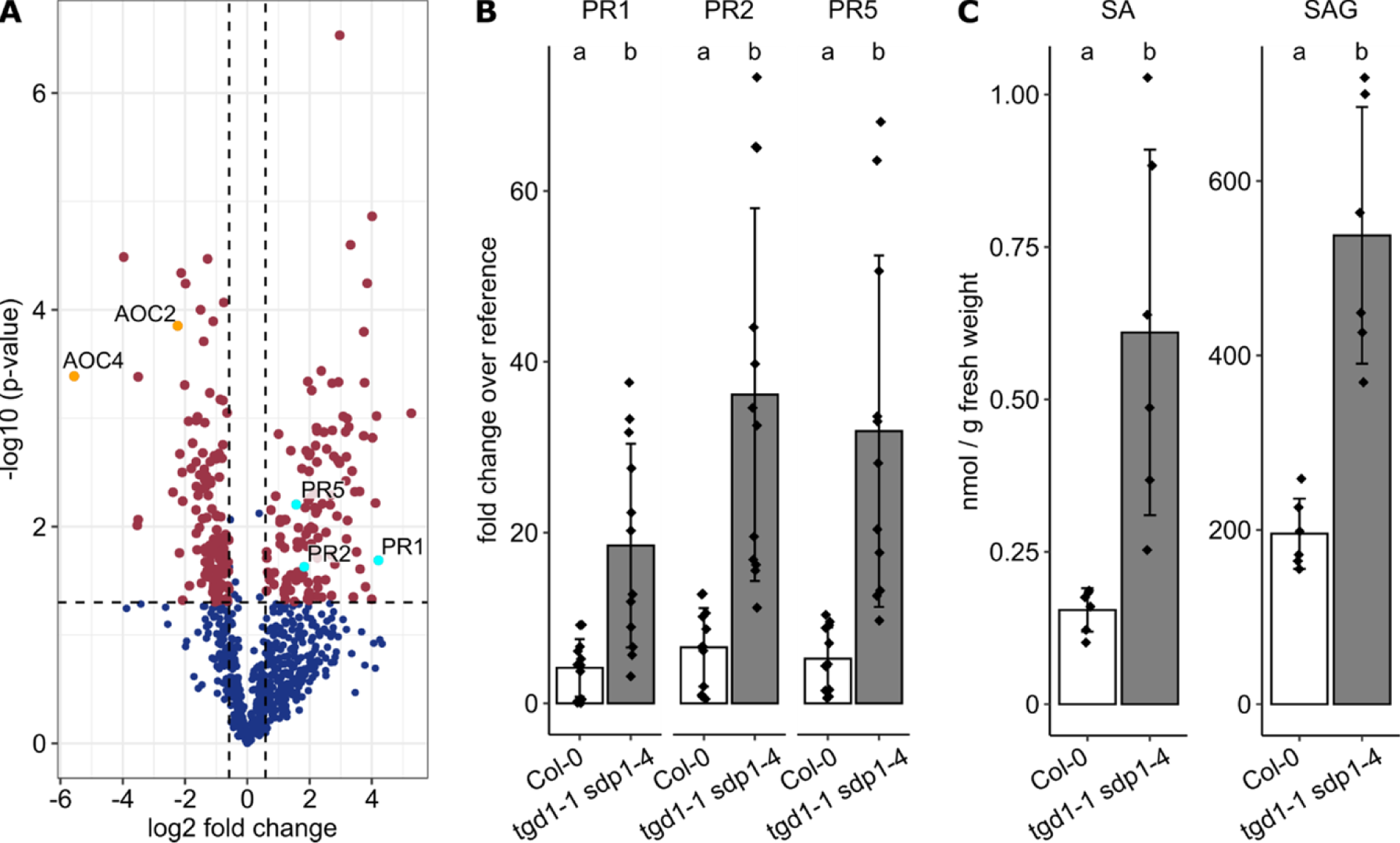
Differences in the proteome of Arabidopsis Col-0 compared to the double mutant *tgd1-1 sdp1-4*. The proteome of Col-0 and *tgd1-1 sdp1-4* total protein fractions of non-stressed plants was analyzed. Changes in protein abundance are visualized in a volcano plot, displaying proteins accumulated (upper right) or depleted (upper left) in the double mutant (A). Proteins were only included in the analysis if they were present in all replicate samples of at least one line. Vertical lines indicate 1.5-fold enrichment or depletion, and the horizontal line indicates a *p*-value of 0.05. Proteins further discussed in the text are marked: the glutathione S-transferase GSTF7 and the PR proteins PR1, PR2 and PR5 accumulate in the mutant (cyan dots), while the allene oxide cyclases AOC2 and AOC4 are depleted (orange dots). *PR* gene expression was further analyzed in leaves of Col-0 or *tgd1-1 sdp1-4* and expression levels were calculated relative to the reference gene *PTB1* (*AT3G01150)* (B). Leaves of both plant lines were also analyzed by UPLC-nanoESI-MS/MS for their salicylic acid (SA) and SA glucoside (SAG) content (C). p-values in (A) were calculated by Student’s t-test. Values are shown as mean ± standard deviation in (B) and (C). Statistical analysis in (B) and (C) was carried out with the Wilcoxon-Mann-Whitney-Test, using Holm-Bonferroni correction for multiple comparisons. Statistical differences with *p*<0.05 are indicated by different letters. n≥3 biological replicates in (A), n=6 biological replicates in (B) and (C). For (B) two independent technical replicates of each biological replicate were measured.

When analyzed with the STRING web tool, two protein-interaction networks among the downregulated proteins were most striking (Suppl. Figure S8). That is, several components of the light harvesting complexes (among others LHCA3, LHCB3) and the photosynthetic electron transfer chain (e.g. PSBA, PSBE, PETA, PETD, PSAE-2) were decreased in the double mutant. The same was true for several ribosomal proteins of both cytosolic and plastidial ribosomes. Among the upregulated proteins in the double mutant, enzymes of various metabolic pathways could be found, including four glutamine synthetases (GLN1;1, GLN1;2, GLN1;3, GS2) and three proteins of the glycine cleavage system (GLDT, GLDP1, GDC-H1) that plays a role in photorespiration (Bauwe et al., 2010).

Among the less abundant proteins in the *tgd1-1 sdp1-4* mutant proteome, we observed two allene oxide cyclase proteins, AOC2 and AOC4, which catalyze the cyclisation step in the biosynthesis of jasmonic acid (JA, Figure 5A). In contrast, the SA-related proteins PR2, PR3 and PR5 were upregulated and this upregulation in comparison to the wild type was also observed under stress conditions (Suppl. Figure S9). Following up on this possible change in SA-related signaling, we tested the gene expression of *PR1*, *PR2* and *PR5* in non-stressed leaves of Col-0 and *tgd1-1 sdp1-4* by qPCR and observed increased transcript levels in the mutant (Figure 5B). Finally, we also measured phytohormone levels in leaves. In line with the observed changes of *PR* gene expression and protein abundance, the base level of SA in non-stressed plants is increased in *tgd1-1 sdp1-4,* as is the amount of its glycosylated derivative, salicylic acid glucoside (SAG; Figure 5C). Due to the low amount of JA and JA-derivatives in non-stressed plants, we were not able to determine if their basal concentrations were also affected, as the lower protein amounts of AOC2 and AOC4 suggested.

### Survey of proteins enriched at Arabidopsis leaf LDs reveals LD localization of LDNP and CB5-E

An important aim of this work was to identify proteins so far unknown to localize to LDs in leaves, since our understanding of LD biology hinges on the understanding of its associated proteins and their functions. We therefore obtained an enriched LD fraction from leaves of *tgd1-1 sdp1-4* mutant Arabidopsis plants and investigated the proteome using quantitative label-free proteomics. First, to evaluate the success of LD enrichment, we combined the riBAQ values of all known detected LD proteins (Gidda et al., 2016; Brocard et al., 2017; Pyc et al., 2017; Kretzschmar et al., 2018; Fernández-Santos et al., 2020; Kretzschmar et al., 2020; Doner et al., 2021; Ge et al., 2022; Li et al., 2022) revealing a strong enrichment of this combined abundance by a factor of 65 to 775 in the LD fractions of the different treatment conditions (Suppl. Figure S10). We then tested if other organelles co-enrich with LDs. Using the plant proteome database (PPDB; http://ppdb.tc.cornell.edu/; Sun et al., 2009), the abundance of all proteins with an assigned unique subcellular localization was combined. The ER and plastoglobule proteomes were most prominently co-enriched by factors of 4.8 to 45 and 4.9 to 33, respectively, which might be a reflection of LD biogenesis at the ER and the similarities in the density of cytosolic LDs and chloroplastic plastoglobuli. All other subcellular structures were either not strongly enriched or significantly depleted in the LD-enriched fractions (Suppl. Figure S10).

In order to identify potential new LD proteins, we then calculated the enrichment and its statistical significance for individual proteins (Suppl. Dataset S17). By combining all datasets from the different treatments, we were able to identify proteins that consistently showed a higher accumulation in the LD-enriched fractions (Figure 6). In total, 553 proteins significantly accumulated in the LD-enriched fractions, 102 of which had an enrichment factor of more than 16 (Figure 6B). Among these were several known LD proteins, most prominently CLO3, LDAP3, and LDIP (Shimada et al., 2014a; Gidda et al., 2016; Pyc et al., 2017). Another protein with a high enrichment value was SEC61γ, a subunit of the SEC61 translocon previously described at ER-LD contact sites (Kretzschmar et al., 2020). A high LD enrichment was also observed for a protein (AT5G04830) that is annotated at The Arabidopsis Information Resource (TAIR; Berardini et al., 2015) as NUCLEAR TRANSPORT FACTOR 2 (NTF2) family member but has not been functionally characterized. Additional proteins that were studied further were selected based on their enrichment value, predicted transmembrane (TMD) regions, and/or possible LD-related functions. Among these were two cytochrome *b*_5_ proteins, CB5-D (AT5G48810) and CB5-E (AT5G53560), that were both highly enriched in the LD fraction (Figure 6B) and might serve in electron transfer during possible lipid oxidation reactions at LDs. We also examined the intrinsically disordered protein EARLY RESPONSIVE TO DEHYDRATION 14 (AT1G76180), which is proposed to act as chaperone in stressed plants (Kovacs et al., 2008; Szalainé Ágoston et al., 2011), as well as six other so far uncharacterized proteins, i.e., AT1G65270, AT1G72170, AT3G18430, AT4G12590, AT4G16450 and AT5G01750, some of which are annotated (at TAIR) to contain domains of unknown function (AT1G72170, AT4G12590, AT5G01750) and some are annotated to be part of membrane protein complexes (AT1G65270, AT1G72170, AT4G12590). Further, we focused on two other proteins, AT3G18430 and AT4G16450, that are annotated as a calcium-binding EF-hand family protein and NADH-ubiquinone oxidoreductase, respectively, and therefore drew our attention as being potentially involved in signaling or redox processes at the LD.

**Figure 6:**
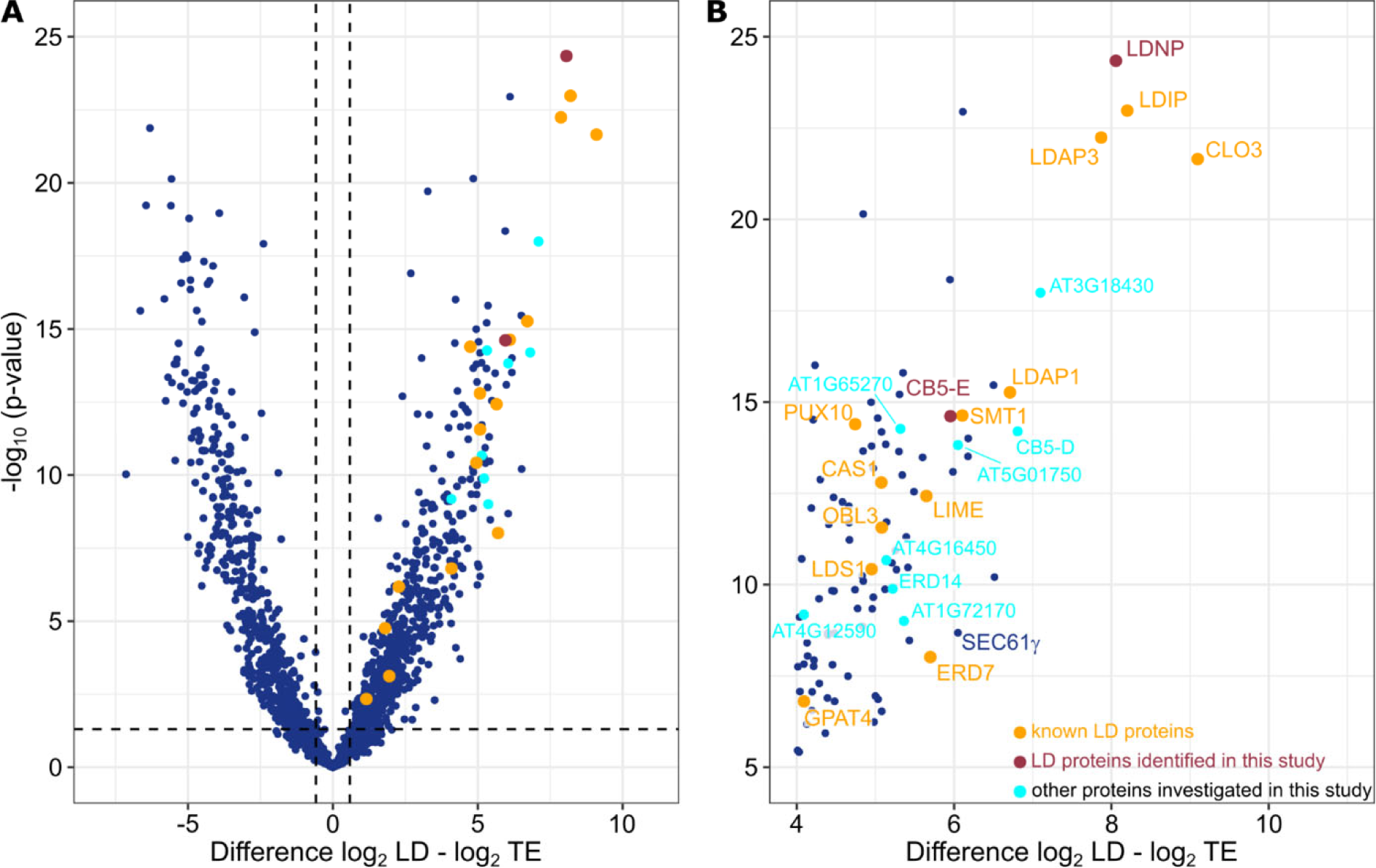
Enrichment analysis of proteins in the LD-enriched fractions prepared from Arabidopsis leaves. LDs were enriched from leaves of Arabidopsis *tgd1-1 sdp1-4* plants that were either untreated or subjected to different stresses. Subsequently, the proteome of the LD-enriched fractions and the corresponding total leaf protein extract was measured. Of the detected proteins, a volcano plot was created, plotting the enrichment of each protein in the LD-fraction against its respectively calculated *p*-value (A). Treatments were combined, however, proteins were only included in the analysis if they were identified by at least two peptides and were present in at least three replicates in one of the sample types. Proteins significantly enriched in the LD-fraction cluster in the upper right corner and this section of the volcano plot is depicted enlarged in (B). Previously known LD proteins are marked in orange; proteins investigated in this study that did or did not localize to LDs are highlighted in red and cyan, respectively. Known and new LD proteins are labeled, in addition the protein SEC61γ is indicated. *P*-values were calculated by Student’s t-test. Vertical lines indicate 1.5-fold enrichment or depletion, while the horizontal line indicates a significance of *p* < 0.05. LD, lipid droplet; TE, total protein extract; CAS1, CYCLOARTENOL SYNTHASE 1; CB5-D/E, CYTOCHROME B5 ISOFORM D/E; CLO3, CALEOSIN 3; ERD7/14, EARLY-RESPONSIVE TO DEHYDRATION 7/14; GPAT4, GLYCEROL-3-PHOSPHATE ACYLTRANSFERASE 4; LDAP1/3, LD-ASSOCIATED PROTEIN 1/3; LDIP, LDAP INTERACTING PROTEIN; LDNP, LD-LOCALISED NTF2 FAMILY PROTEIN; LDS1, LIPID DROPLETS AND STOMATA 1; LIME, LD-associated methyltransferase; OBL3, OIL BODY LIPASE 3; PUX10, PLANT UBX DOMAIN CONTAINING PROTEIN 10; SEC61γ, SEC61 GAMMA; SMT1, STEROL METHYLTRANSFERASE 1.

We analyzed the subcellular localization of all the above-mentioned candidate LD proteins by transient expression in *Nicotiana benthamiana* leaves, which is a well-established model plant system for protein localization (Sparkes et al., 2006), including LD proteins (Kretzschmar et al., 2020; Doner et al., 2021; Pyc et al., 2021; Krawczyk et al., 2022b). Candidates were expressed with an N- or C-terminal mCherry fluorescent tag and subcellular localization was analyzed by confocal laser-scanning microscopy (CLSM). To assess possible LD co-localization, LDs were stained with the neutral lipid-specific stain BODIPY 493/503 (Listenberger and Brown, 2007). Furthermore, selected candidate LD proteins were co-expressed with the mouse (*Mus musculus*) enzyme DIACYLGLYCEROL ACYLTRANSFERASE 2 (MmDGAT2), which enhances LD proliferation in plant cells (Cai et al., 2019). As shown in Figure 7 the NTF2 protein family member often localized to the surface of BODIPY-stained LDs, particularly in cells co-expressing MmDGAT2 (Figure 7A, B). We therefore termed this protein <u>LD</u>-LOCALISED <u>N</u>TF2 FAMILY <u>P</u>ROTEIN (LDNP). For CB5-E, we observed two distinct subcellular localizations depending on the position of the appended mCherry: the C-terminal mCherry-tagged CB5-E (CB5-E-mCherry) localized to reticular structures, consistent with the ER (Figure 7C), while N-terminal mCherry-tagged CB5-E (mCherry-CB5-E) displayed mostly LD localization (Figure 7D). Given that bioinformatic tools predict a single TMD near the C-terminus of CB5-E (Suppl. Figure S11), this region might be less accessible with a C-terminal-appended mCherry and therefore resulting in the observed differences in CB5-E localization. Overall, the other eight candidate proteins appended to mCherry at their N and/or C-termini did not localize to BODIPY-stained LDs, although CB5-D appeared to partially localize to LDs and some proteins (e.g., AT5G01750 and AT1G76180 [ERD14]) displayed a distinct reticular, ER-like fluorescence patterns that were often closely associated with LDs (Suppl. Figure S12), which may be notable given the role of the ER in LD biogenesis (Guzha et al., 2023). Indeed, the apparent association of ERD14 in reticulum-like structures associated with LDs was more pronounced upon co-expression with MmDGAT2 (Suppl. Figure S12), suggesting a possible role for ERD14 at the ER during LD proliferation.

**Figure 7:**
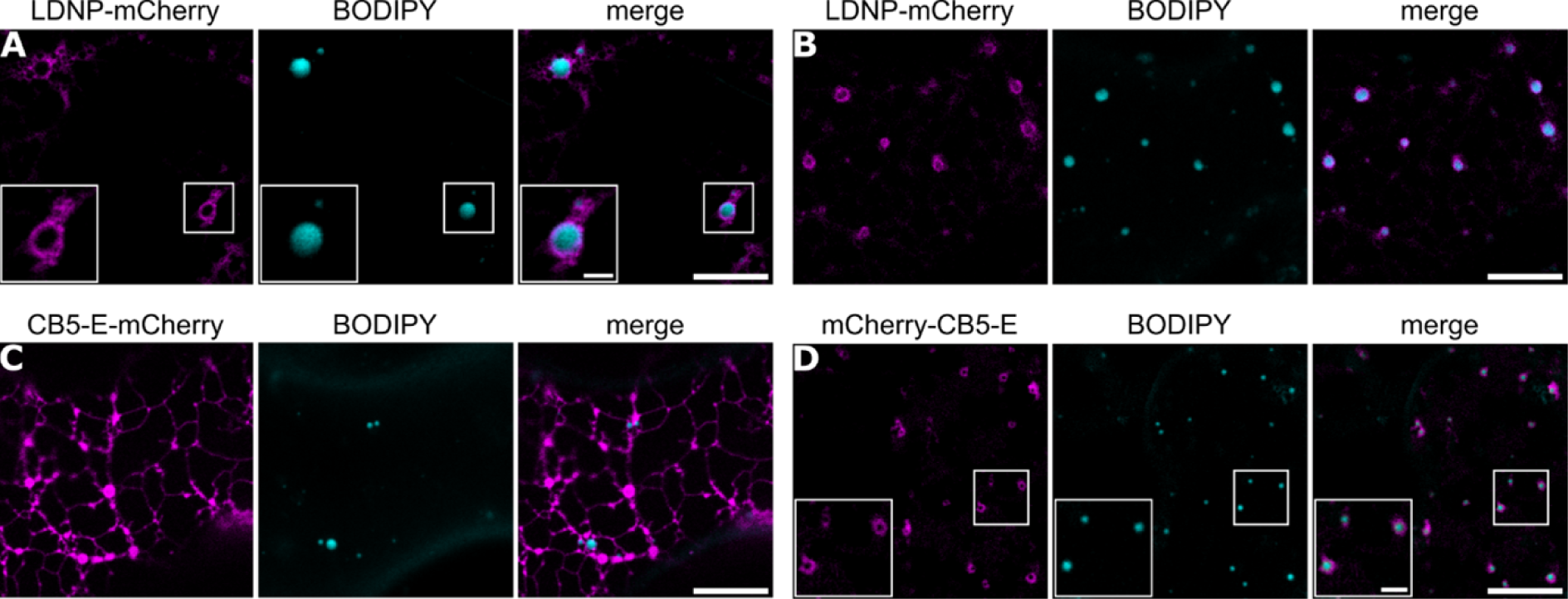
LDNP and CB5-E loclaize to LDs in *Nicotiana benthamiana* leaf cells. Subcellular localization studies of N- and/or C-terminal mCherry-tagged LDNP and CB5-E were carried out by transient expression in *N. benthamiana* leaves. Proteins were fused to an mCherry-tag (magenta channel) and LDs were stained with BODIPY 493/503 (cyan channel). Shown are also the corresponding merged images; boxes in (A, D) highlight regions of the cell shown with higher magnification in the insets. LDNP-mCherry localization to BODIPY-stained LDs was observed when the protein was expressed alone (A) or when co-expressed with MmDGAT2 (B), which causes a proliferation of LDs in plant cells (Cai et al., 2019). CB5-E appended at its C-terminus to mCherry did not localize to LDs (C), however, N-terminal mCherry-tagged CB5-E localized to LDs (D). Bars = 10 µm (2 µm in insets).

### The Arabidopsis leaf LD proteome responds to environmental stresses

Based on an examination of the LD proteome in leaves of *tgd1-1 sdp1-4* mutant plants without stress treatment, CLO3 and LDAP3 were the most abundant proteins (Table 1) as they combined abundance amounts to around 80% of total LD protein. Additional proteins detected in the LDs of leaves without treatment included LDAP1 and LDAP2, their interacting protein LDIP, and the protein ERD7 (Gidda et al., 2016; Pyc et al., 2017; Doner et al., 2021). Several other proteins connected to metabolism (i.e., GPAT4, STEROL METHYLTRANSFERASE 1, CYCLOARTENOL SYNTHASE 1, OIL BODY LIPASE 3, LD-ASSOCIATED LIPASE 2, α-DOX1, LD-ASSOCIATED METHYLTRANSFERASE 1 or 2) and protein degradation (PLANT UBX DOMAIN-CONTAINING PROTEIN) were also found in smaller amounts. In addition, LDNP and CB5-E described above were detected (Table 1).

**Table 1:**
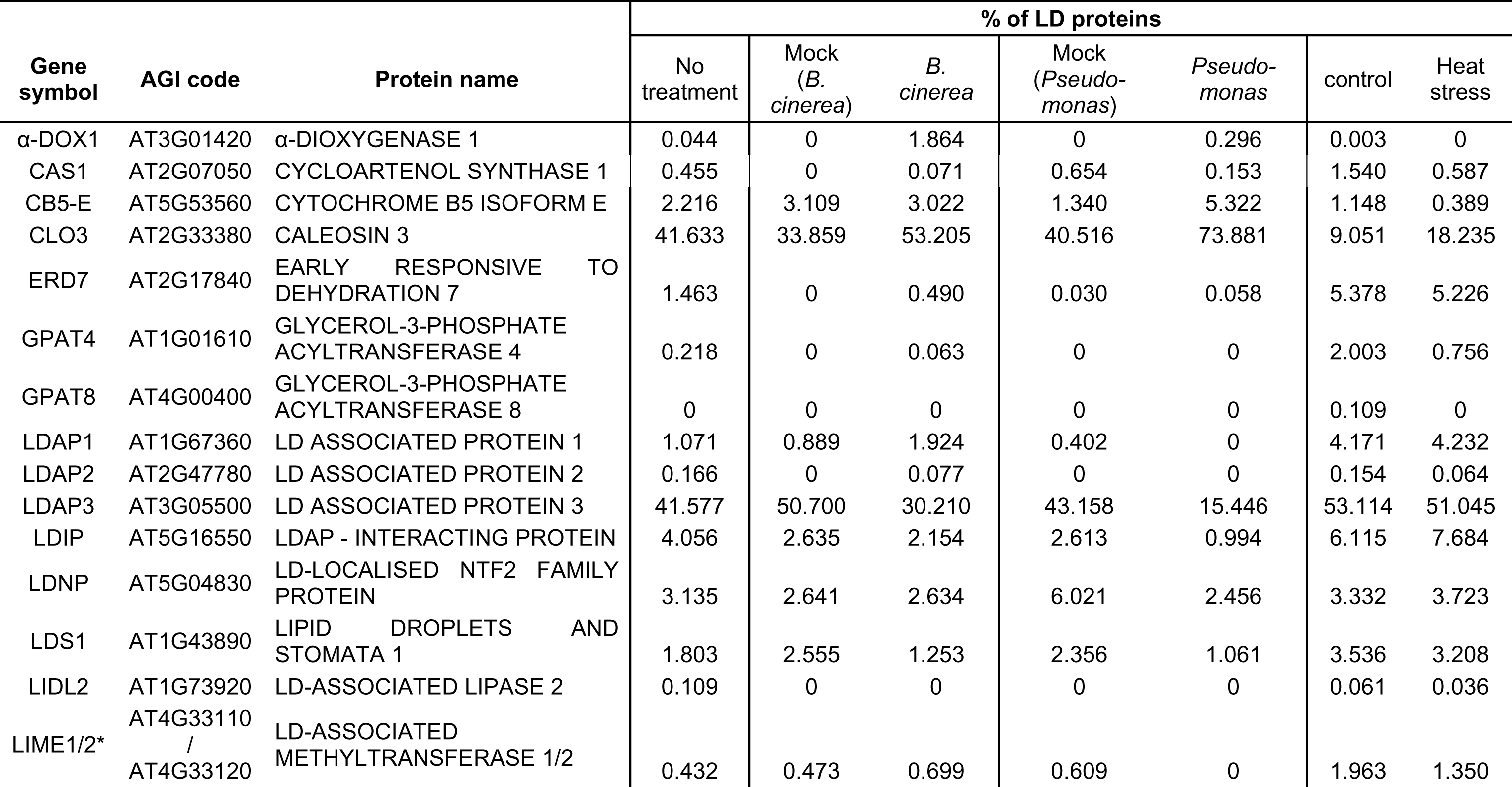
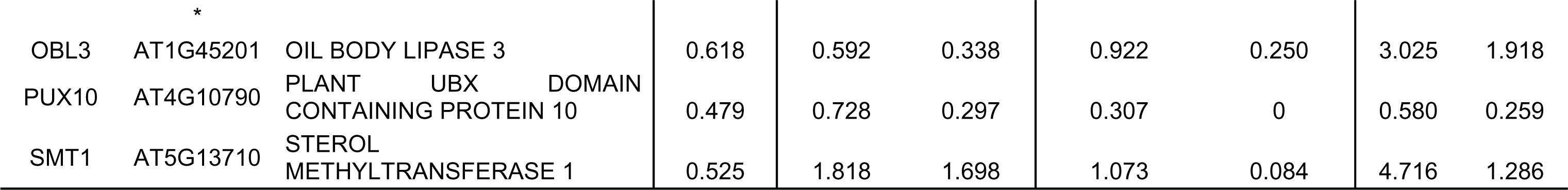
Composition of LD proteome in LD-enriched fractions of leaves from Arabidopsis *tgd1-1 sdp1-4*. LDs were isolated from Arabidopsis leaves *of tgd1-1 sdp1-4* in non-stressed conditions and after different stresses. Stress treatments of leaves included infection with *Botrytis cinerea* (*B. cinerea*) or *Pseudomonas syringae* pv. *tomato* DC3000 *ΔavrPto ΔavrPtoB* (*Pseudomonas*), or heat stress for 24 hours at 37°C. For each stress, a mock or control treatment was performed. The relative contribution of individual LD proteins to the total LD protein abundance (riBAQ values) was calculated for each treatment separately. * denotes proteins that could not be identified unequivocally.

We did not observe any additional LD proteins specific to the different stress treatments. However, we were able to follow the changes in the LD proteome composition in response to the individual stresses (Figure 8). Overall, the combined abundance of LD proteins tended to increase after stress, especially after infections (Figure 8A). However, increased abundance was not evenly distributed across all LD proteins, so that the relative proportions of detected LD proteins in the known LD proteome changed (Figure 8B, Table 1). For example, the proportion of CLO3 increased after all three treatments in contrast to LDAP3, whose relative contribution either decreased or did not change (Figure 8C). Among the low-abundant leaf LD proteins, α-DOX1 strongly accumulated at LDs after both pathogen treatments, which fits with the previously described functional interaction between CLO3 and α-DOX1 (Shimada et al., 2014b). Interestingly, α-DOX1 did not increase in response to heat treatment (Figure 8C), pointing to a more specific role in defense against pathogens. For CB5-E, we also observed its accumulation specifically after pathogen infection, although on a smaller scale than for α-DOX1 (Figure 8C). Furthermore, LD-ASSOCIATED METHYLTRANSFERASE 1 or 2 (the distinct isoform could not be resolved based on the analysis) increased under *B. cinerea* but not *Pseudomonas* infection. Most other proteins fluctuated in abundance but showed no clear trend with the possible exception of PUX10, which was depleted after all the treatments (Figure 8C).

**Figure 8:**
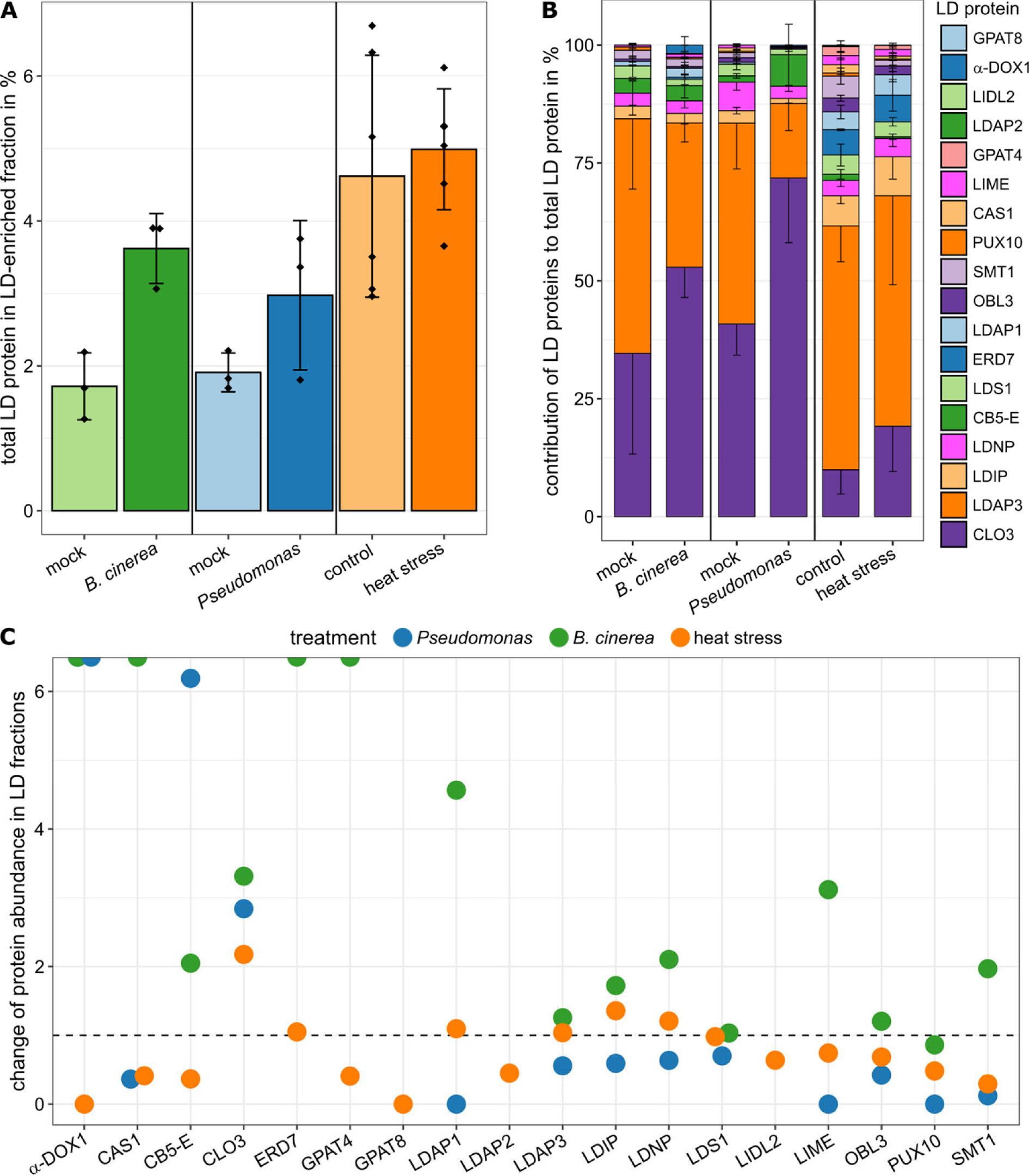
Changes in the known LD proteome of leaves after stress treatments. The protein abundance (riBAQ values) of known LD proteins in the LD-enriched fraction isolated from Arabidopsis leaves was followed in reaction to infection with *Botrytis cinerea* (*B. cinerea*), or *Pto* DC3000 *ΔavrPto/ΔavrPtoB* (*Pseudomonas*), or heat stress for 24 h at 37°C. The total LD protein abundance was calculated by summing up riBAQ values of all LD proteins for each treatment (A). In addition, the relative contribution of all detected LD proteins to the total LD protein abundance was calculated (B). Stacked bar plots show the relative proportion of individual LD proteins in the order displayed in the legend, i.e. percentage of GPAT8 at the top and percentage of CLO3 at the bottom. The changes in abundance of individual proteins was followed by calculating the ratio of their riBAQ values in LD fractions of stressed plants relative to their riBAQ values in respective control treatments (C). Values above and below 1 indicate enrichment or depletion upon individual stresses, respectively. The horizontal line highlights ratios of 1. Proteins were only included if they were identified by at least two peptides and were detected in at least three replicates of at least one sample type. n≥3 replicates per treatment. α-DOX1, α-DIOXYGENASE 1; CAS1, CYCLOARTENOL SYNTHASE 1; CB5-E, CYTOCHROME B5 ISOFORM E; CLO3, CALEOSIN 3; ERD7, EARLY-RESPONSIVE TO DEHYDRATION 7; GPAT4/8, GLYCEROL-3-PHOSPHATE ACYLTRANSFERASE 4/8; LDAP1/2/3, LD-ASSOCIATED PROTEIN 1/2/3; LDIP, LDAP INTERACTING PROTEIN; LDNP, LD-LOCALISED NTF2 FAMILY PROTEIN; LDS1, LIPID DROPLETS AND STOMATA 1; LIDL2, LIDL2, LD-ASSOCIATED LIPASE 2; LIME, LD-ASSOCIATED METHYLTRANSFERASE 1/2 (ambiguously identified); OBL3, OIL BODY LIPASE 3; PUX10, PLANT UBX DOMAIN CONTAINING PROTEIN; SEC61γ, SEC61 GAMMA; SMT1, STEROL METHYLTRANSFERASE 1.

## Discussion

### Biotic and abiotic stresses alter lipid homeostasis in Arabidopsis leaves

Higher temperatures lead to increased membrane fluidity so that membrane lipids are usually remodeled and acyl chains with three double bonds are replaced by more saturated acyl chains (Falcone et al., 2004; Higashi and Saito, 2019; Yu et al., 2021). Such a remodeling was also observed in our study with presumably 16:0 and 18:0-containing lipid species increasing especially in PC, and 18:3 in MGDG and DGDG being replaced by 18:2 and 18:1 (Figure 2). TAG acts as sink for the released acyl chains (Mueller et al., 2017), which is reflected in the increase of 54:8 and 54:9-TAG species after heat stress that we and others observed (Higashi et al., 2015; Mueller et al., 2015; Mueller et al., 2017). Interestingly, the amount of SEs decreased (Figure 1), corroborating previous results (Shiva et al., 2020). Furthermore, other sterol derivatives like sterol glycosides and acylated sterol glycosides have been reported to increase in response to heat stress in Arabidopsis leaves and tobacco pollen tubes (Shiva et al., 2020; Krawczyk et al., 2022a). This indicates that LDs cannot only serve as a sink for acyl chains stored as TAG, but at the same time might be a source for free sterols or sterol derivatives derived from LD-stored SEs. These sterols could help to stabilize membranes under heat stress (Dufourc, 2008). Regarding the potential dual role of LDs as a metabolic sink and source, it is unclear if SEs and TAGs in plant cells are both present in mixed-compound LDs or form distinct LD subpopulations, as has been reported for LDs in some animal cells (Khor et al., 2014).

Unlike lipidomic changes after heat stress, less is known about the impact of pathogen infection. Infection of leaves with the avirulent *Pseudomonas* strain *Pseudomonas syringae* pv. *tomato* DC3000 *avrRpm1* causes an increase in TAG levels (Schieferle et al., 2021) and leads to increased numbers of leaf LDs (Fernández-Santos et al., 2020) within one day after infection. Using another avirulent *Pseudomonas* strain (*Pseudomonas syringae* pv. *tomato* DC3000 *ΔavrPto/ΔavrPtoB*) and measuring neutral lipids after symptom development, we observed a similar trend to increased TAG levels (Figure 1), which was mainly driven by the TAG species 54:8 and 54:9 (Suppl. Datasets S4, S6). Fungal infection of Arabidopsis with *B. cinerea* in leaves (Figure 1) and *Verticillium longisporum* in roots (Schieferle et al., 2021) are also accompanied by a trend towards increased TAG levels. Interestingly, in case of *V. longisporum*, TAG accumulation in leaves was proposed as a systemic effect initiated by the infected roots (Schieferle et al., 2021). It thus seems that increased TAG levels are a general plant response to infection. However, the reasons for that are not clear yet. With regard to sterol lipids, the most striking change after infection is the increase of stigmasterol at the expense of β-sitosterol (Figure 3). This effect on free sterols has been described previously (Griebel and Zeier, 2010; Wang et al., 2012), however, descriptions of its role in the plant-pathogen interaction are conflicting. In Arabidopsis, the conversion of β-sitosterol to stigmasterol is catalyzed by the cytochrome P450 enzyme AtCYP710A1 (Morikawa et al., 2006), and T-DNA mutants of *CYP710A1* with decreased stigmasterol levels were reported to be more resistant to infection with *P. syringae* pv. *maculicola* (Griebel and Zeier, 2010). Later reports on another *cyp710a1* mutant described the opposite effect though, as the mutant was more susceptible to a variety of *P. syringae* strains (Wang et al., 2012).

The substantial changes observed in the lipidome especially under heat stress are not reflected in the abundance of proteins involved in lipid metabolism. This could be due to low coverage of such proteins, as only around 20% of the proteins were detected in our data (Suppl. Dataset S12). Previous reports on heat-stressed leaves indicated regulation of lipid-related genes on the transcriptional level (Higashi et al., 2015). However, in a study on the transcript changes in pollen tubes under heat stress (Krawczyk et al., 2022a), only few changes were observed. As of this it can be speculated that lipid remodeling is also controlled by post-translational modifications or other factors.

### The Arabidopsis cellular proteome is readjusted under stress

One common theme under stress was the reduction in the levels of photosynthesis-related proteins. This could be a protection mechanism to reduce photosynthesis rates given that chlorophyll fluorescence studies have shown reduced quantum efficiency in response to infection (Bonfig et al., 2006; Pavicic et al., 2021) or heat stress (Kim and Portis, 2005; Salvucci, 2007). On the other hand, oxidative stress in the plastids could also lead to an increased damage and degradation of photosynthesis-related proteins. In addition, heat stress is especially linked to decreases in RuBisCO activity (Kobza and Edwards, 1987; Kim and Portis, 2005; Salvucci, 2007), consistent with our observation of RuBisCO subunits among the depleted proteins (Suppl. Figure S5). Furthermore, for infection, several transcriptomics studies have shown that reduced gene expression of photosynthetic genes is a general response to a plethora of pathogens, possibly to allow for the upregulation of defense response pathways (Bilgin et al., 2010; Jiang et al., 2017).

Proteins that were upregulated were more diverse between the treatments, conferring specificity to the plant’s reaction. In the proteome of heat-stressed plants this resulted in an accumulation of heat shock proteins (HSPs; Figure 4), as expected. HSPs prevent the formation of protein aggregates and assist protein folding and protein transport across membranes (Lin et al., 2001; Rosenzweig et al., 2019). Additionally, several chloroplast chaperonin subunits were upregulated in response to heat stress, including CPN60A1 and CPN60B1 of chaperonin 60, which interacts in the assembly of RuBisCO (Hemmingsen et al., 1988; Ishikawa et al., 2003).

In the proteome of infected leaves, several metabolic proteins were upregulated, including glutathione S-transferases (GSTs) after both infections (Suppl. Figure 5). Different GSTs were previously reported to change in abundance in a 2D-proteomics study of Arabidopsis infected with *Alternaria brassicicola* (Mukherjee et al., 2010). GSTs thus may form part of a general defense responses, although the functional role of various GST proteins are likely different. GSTF6, for example, has been implicated in the biosynthesis of camalexin (Su et al., 2011), a phytoalexin with antifungal properties against some *B. cinerea* strains (Kliebenstein et al., 2005). GSTF2, on the other hand, was suggested to bind small molecules, including antimicrobial compounds like camalexin, and to transport them within the cell to their site of action (Dixon et al., 2010; Ahmad et al., 2017). Further metabolic pathways upregulated after *B. cinerea* infection were tryptophan biosynthesis and cyanide detoxification. Both might be connected to synthesis of camalexin, as it derives from tryptophan, and in camalexin biosynthesis, cyanide is released (Böttcher et al., 2009). Interestingly, the phospholipase PLA2A, which was upregulated after *B. cinerea* infection, has been described to negatively influence plant resistance to *B. cinerea* (La Camera et al., 2005), indicating that metabolic adjustments are influenced by both the plant and the pathogen. Altogether, the rewiring of metabolism appears as a common feature after pathogen infection, and the selectivity of altered metabolic pathways conveys the distinct responses towards specific pathogens.

### Effects of the *tgd1-1 sdp1-4* mutations in Arabidopsis are not limited to lipid metabolism

We used the Arabidopsis double mutant *tgd1-1 sdp1-4* as a tool for LD isolation, and we also included it in our lipidomic and proteomic analyses. The original characterization of the double mutant had highlighted its strong increase in TAG levels in leaves (Fan et al., 2014) and we confirmed this increase in our lipidomics measurements (Suppl. Figure 6). While the increase of TAG is caused by both the mutations in *SDP1* and *TGD1*, the changes in the membrane lipidome are likely mostly caused by the mutation of TGD1 alone. TGD1 is involved in the import of membrane lipid precursors into plastids, and its loss in the *tgd1-1* mutant leads to a moderate reduction of thylakoid galactolipids and an altered FA profile of galactolipids, PC, PE and PA (Xu et al., 2003; Fan et al., 2014). We found similar alterations not only in PG that predominantly occurs in plastids (Joyard et al., 2004) but also in phosphatidylinositol and phosphatidylserine, indicating that also non-plastidial lipid metabolism is strongly affected.

Regarding the proteome of the mutant, it is notable that we detected LDAP3 as the most abundant protein of the LDAP family (Figure 8B, Table 1). In previous proteomic studies of senescent or drought-stressed leaves of Arabidopsis Col-0, LDAP1 was the major LDAP protein (Brocard et al., 2017; Doner et al., 2021). A possible explanation could be that expression of *LDAP1* in contrast to *LDAP3* is upregulated in response to drought stress (Wilkins et al., 2010) and during leaf senescence (Schmid et al., 2005). Apart from the differences between LDAP1 and LDAP3, the composition of the LD proteome is overall quite similar to previously-published LD proteomes in senescent and drought-stressed Col-0 leaves (Brocard et al., 2017; Doner et al., 2021). As well as in these other proteomic studies, CLO3 was a dominant LD protein and, along with the LDAP proteins, comprised up to more than 90% of the LD proteome. Other less abundant LD proteins were also found, including LDIP and α-DOX1, with relative contributions to the total LD proteome that did not exceed 2% (Brocard et al., 2017; Doner et al., 2021) and thus similar to our observations in *tgd1-1 sdp1-4* (Table 1).

Besides the proteome of LD-enriched fractions, we also analyzed total protein fractions from leaves of *tgd1-1 sdp1-4*, which we compared to total protein fractions from Arabidopsis wild type. This comparison revealed the upregulation of the defense-related proteins PR1, PR2 and PR5 and the concomitant increase in SA levels in *tgd1-1 sdp1-4* (Figure 5). Neither TGD1 nor SDP1 have been linked to SA synthesis in Arabidopsis (Rekhter et al., 2019). However, previous studies reported that interference with plastid lipid metabolism impacts endogenous levels of jasmonates (Lin et al., 2016). Knockout of a gene encoding a chloroplast outer envelope protein that synthesizes DGDG caused the concomitant increase of JA and JA-Ile (Lin et al., 2016). Assuming that the *tgd1* mutation might also affect phytohormones that derive from plastids, the observed differing levels of SA in *tgd1-1 sdp1-4* could then be a secondary effect on the crosstalk between the phytohormones salicylic acid and jasmonates (Spoel et al., 2003; Pieterse et al., 2012). In this context, it is interesting to note that reduced growth is a phenotype of both *tgd1-1* and *tgd1-1 sdp1-4* (Xu et al., 2005; Fan et al., 2014). Given the differences in TAG accumulation between the single and the double mutant (Fan et al., 2014), reduced growth is probably not caused by the hyperaccumulation of TAGs, instead, autoimmune reactions might contribute to this growth difference. If that is the case, connections between autoimmunity and mutations designed to increase TAG levels should be considered in any future biotechnological approaches aimed at the hyperaccumulation of TAG in vegetative tissues.

### Observed LD-localization of Arabidopsis LDNP and CB5-E indicates new LD functions

While the number of known LD-associated plant protein families has steadily increased in recent years (Fernández-Santos et al., 2020; Kretzschmar et al., 2020; Doner et al., 2021; Ge et al., 2022; Krawczyk et al., 2022b; Li et al., 2022), the number is still only roughly two dozen (Guzha et al., 2023). These studies were often followed up by the functional characterizations of these proteins, greatly increasing our understanding of LDs (Shimada et al., 2014a; Gidda et al., 2016; Deruyffelaere et al., 2018; Kretzschmar et al., 2018; Pyc et al., 2021; Krawczyk et al., 2022b). Here, we identified two additional proteins with the ability to localize to LDs in plant leaves, CB5-E and LDNP (Figure 7). Of these, the LD localization of LDNP became more distinct upon LD proliferation due to the co-expression with MmDGAT2, which could indicate that LDNP requires a certain threshold number of LDs to localize to LDs. CB5-E and LDNP were also shown previously to be enriched in LD-fractions from Arabidopsis seedlings and drought-stressed leaves, but neither protein was investigated in regard to LD targeting (Kretzschmar et al., 2018; Kretzschmar et al., 2020; Doner et al., 2021). CB5-E was originally described as cytochrome *b*_5_ protein able to accept electrons from the NADH-dependent cytochrome *b*_5_ reductase CBR (Fukuchi-Mizutani et al., 1999). CB5-E is part of a five-member protein family in Arabidopsis, and its homolog CB5-D was also enriched in our LD fractions (Suppl. Dataset S17). Arabidopsis CBR and CB5 proteins are involved in desaturation reactions (Kumar et al., 2006; Kumar et al., 2012). In microsomal fractions of castor bean, desaturation and hydroxylation reactions in the synthesis of ricinoleic acid depended on cytochrome *b*_5_ proteins (Smith et al., 1992). CB5-D has also been previously localized to the ER (Maggio et al., 2007). We could indeed observe ER localization for CB5-D and CB5-E. However, CB5-E also localized predominantly to LDs in cells co-expressing MmDGAT2 (Figure 7). This raises the question if particularly CB5-E could be involved in additional redox reactions at the LD surface during LD proliferation.

In comparison to CB5-E, much less is known about the function(s) of LDNP. Expression of *LDNP* did not show specific tissue or developmental preferences (Klepikova et al., 2016) and the protein is also found in LD fractions of 24 h – 60 h old seedlings and drought-stressed leaves of Arabidopsis (Kretzschmar et al., 2020; Doner et al., 2021). LDNP was classified as a NTF2 family protein, probably based on a predicted protein fold, which is thought to form a cone-like shape with internal cavity (Eberhardt et al., 2013). This protein structure is also found in several other enzymatic and non-enzymatic proteins (Eberhardt et al., 2013), making LDNP an interesting target for further functional characterization.

### Changes in the Arabidopsis LD proteome establish LDs as an additional player in stress responses

We analyzed the LD proteome in leaves of the *tgd1-1 sdp1-4* double mutant after different stresses and observed that it was altered dynamically (Figure 8B). When analyzing these changes, it has to be considered that there are significant differences in the overall proteome of leaf total protein fractions between the double mutant and the wild type. The observed alterations of the LD proteome of *tgd1-1 sdp1-4* therefore might not be completely the same at leaf LDs in Col-0. Nevertheless, shared upregulated proteins between wild type and the double mutant demonstrate that our treatments induced similar pathways in both Arabidopsis lines. Hence it seems likely that our stress treatments induce analogous reactions in the LD proteome of both the double mutant and the wild type. One example is the increased protein abundances of CLO3 and α-DOX1 that fit well with their previously described roles in pathogen defense in wild-type Arabidopsis plants (Shimada et al., 2014a).

CLO3 and α-DOX1 were among the most strongly responding LD proteins to pathogen treatment, and the protein abundance of CLO3 additionally increased after heat stress (Figure 8). This is in line with reported transcriptome changes of leaves infected with *B. cinerea* and seedlings subjected to heat, salinity and osmotic stress, i.e., *CLO3* expression was induced by all treatments, whereas *α-DOX1* expression was reported as increased during infection and osmotic but not heat treatment (Sham et al., 2015).

How LDs are integrated in plant stress response remains an interesting question. Environmental cues might alter the interaction of LDs with other organelles, e.g. prompting the formation of membrane contact sites for lipid remodeling. A first such contact site of LDs with the plasma membrane has been recently described (Krawczyk et al., 2022b) and although the physiological relevance is unclear as of yet, the interaction of LDs in plant membrane contact sites especially in reaction to stress are an exciting avenue of future LD research.

## Material and Methods

### Plant lines and growth conditions

Lipidomic and proteomic experiments were carried out with Arabidopsis Col-0 and the oil-rich *tgd1-1 sdp1-4* double mutant line (Fan et al., 2014).

Seeds of Arabidopsis lines were surface-sterilized with 6% (w/v) sodium hypochlorite and 0.1% (v/v) Triton X-100 and germinated on half-strength Murashige and Skoog (MS; Duchefa Biochemie, Haarlem, The Netherlands) medium (Murashige and Skoog, 1962) containing 0.8% (w/v) agar. After ten days, seedlings were transferred to soil (Einheitserde SPECIAL Vermehrung, Patzer Erden, Sinntal-Altengronau, Germany) and grown under short-day condition (8 h light/16 h darkness) at 22°C in the light and 18°C in the dark as described previously (Guzha et al., 2022). For lipidomic or proteomic analyses, plants were grown for seven weeks before stress treatment was applied and samples were prepared as described below.

### P. syringae and B. cinerea strains

For plant infections and further proteomic and lipidomic analysis of LDs, the *Pseudomonas* strain *P. syringae* pv. *tomato* (*Pto*) DC3000 *ΔavrPto ΔavrPtoB* and the *Botrytis cinerea* strain B05.10 were used.

Spores of *B. cinerea* were cultured on potato dextrose agar (PDA; Merck KGaA, Darmstadt, Germany) for ten days before conidiospores were harvested by washing them off with ¼ potato dextrose broth (PDB; Merck KGaA, Darmstadt, Germany) and filtering through Miracloth (Merck KGaA, Darmstadt, Germany). Spores were counted with a counting chamber (Fuchs-Rosenthal) and stocks in 15% (v/v) glycerol were stored at –80°C.

### Pathogen treatment for lipidomic and proteomic analysis and subsequent LD-enrichment

For omics-samples of infection treatments, both pathogens were used in spray infections. Infection with *Pto* DC3000 *ΔavrPto ΔavrPtoB* was adapted from (Yao et al., 2013). Briefly, *Pto* DC3000 *ΔavrPto ΔavrPtoB* was cultured in NYG-medium (0.5% [w/v] peptone, 0.3% [w/v] yeast extract, 2% [v/v] glycerol; Merck KGaA, Darmstadt, Germany) with appropriate antibiotics (50 µg/ml rifampicin, 50 µg/ml kanamycin; Duchefa Biochemie, Haarlem, The Netherlands) over night and harvested by centrifugation (1500 x *g*, 20 min, room temperature) at the day of infection. Bacteria were washed once with 10 mM MgCl_2_ and then resuspended in 10 mM MgCl_2_ and 0.02% (v/v) Silwet. Bacterial density was adjusted to an OD_600_ of 1.0 and the bacterial suspension sprayed onto well-watered plants until all leaves were evenly wet. Control plants were sprayed with 10 mM MgCl_2_ and 0.02% (v/v) Silwet. Plants were then covered with plastic hoods to keep them at high humidity.

*B. cinerea* spores from glycerol stocks were diluted to a concentration of 5 x 10^4^ spores/ml in ¼ PDB (Merck KGaA, Darmstadt, Germany) and pre-germinated for 4 hours at room temperature. Plants were sprayed until sprayed droplets ran off the leaves and subsequently kept at humid conditions. Mock-treated plants were treated analogously with ¼ PDB.

Plants were observed until they developed first symptoms to ensure infections were effective and plant proteome alterations had occurred. Sampling was thus carried out 4-5 days after infection by leaf homogenization and LD isolation by ultracentrifugation adapting previous protocols (Kretzschmar et al., 2018). In addition to the pathogen- and mock-treated plants, one proteomic dataset was obtained from plants that were not treated at all.

For LD enrichment used for proteomic analysis, leaves were ground in grinding buffer (50 mM Tris-HCl pH 7.5, 10 mM KCl, 0.4 M sucrose, 200 µM proteinase inhibitor PMSF; Carl Roth GmbH + Co. KG, Karlsruhe, Germany) with sea sand as abrasive agent. Grinding buffer, mortar and pestle were precooled to 0°C and samples were kept between 0 – 4°C during processing. To remove sand and cellular debris, samples were centrifuged for 1 minute at 100 x *g*. An aliquot was taken as total protein extract sample, and proteins were precipitated in 96% ethanol at -20°C. Subsequently, samples were centrifuged at 100,000 x *g* for 35 min in a swing-out rotor. LDs were collected as fat pad from the top of the sample and washed in grinding buffer. After washing, the fat pad was collected and proteins were precipitated with 96% (v/v) ethanol at -20°C.

### Heat stress treatment for lipidomic and proteomic analysis and subsequent LD-enrichment

For heat stress, plants were kept at 37°C for 24 hours, starting with the beginning of the light cycle, while control plants were kept at normal temperatures. For proteomic analysis, LDs were enriched directly after heat stress. To that end, leaves were ground with grinding buffer (10 mM sodium phosphate buffer pH 7.5, 150 mM NaCl, 0.6 M sucrose, 25 mM Lomant’s reagent, 10 mM *N*-ethylmaleimide, 200 µM proteinase inhibitor PMSF; Carl Roth GmbH + Co. KG, Karlsruhe, Germany and Merck KGaA, Darmstadt, Germany). Sea sand was used as abrasive agent and together with cellular debris removed after grinding by centrifugation for 1 minute at 100 x *g*. Total protein extract samples were precipitated from the supernatant at -20°C with 96% ethanol in 10x excess. LDs were enriched by two ultracentrifugation steps at 100,000 x *g* for 60 min in a swing-out rotor. After the first ultracentrifugation, LDs were collected as fat pad and resuspended in washing buffer (10 mM sodium phosphate buffer pH 7.5, 150 mM NaCl, 0.4 M sucrose, 200 µM proteinase inhibitor PMSF, 0.1% [v/v] Tween 20; Carl Roth GmbH + Co. KG, Karlsruhe, Germany). The LD suspension was overlaid with overlay buffer (10 mM sodium phosphate buffer pH 7.5, 150 mM NaCl, 0.2 M sucrose, 200 µM proteinase inhibitor PMSF, 0.1% [v/v] Tween 20; Carl Roth GmbH + Co. KG, Karlsruhe, Germany). After the second ultracentrifugation, LDs were collected as floating fat pad and proteins precipitated at -20°C with 96% ethanol. Before protein precipitation, samples were kept at 0 – 4°C throughout LD enrichment; buffers, mortar and pestle were precooled to the same temperature.

### Lipidomic sample preparation and measurements

Infection with *Pto* DC3000 *ΔavrPto ΔavrPtoB* and *B. cinerea*, and heat stress treatment were carried out as for proteomic samples. Samples were also harvested after the same incubation times and flash-frozen in liquid nitrogen.

Extraction and analysis of triacylglycerol and sterol esters was adapted from (Herrfurth et al., 2021). In brief, frozen leaf material was homogenized with a ball mill, and 500 mg of each sample were extracted by monophasic extraction with 6 ml propan-2-ol:hexane:water (60:26:14, v/v/v) at 60°C. Tripentadecanoin was added as internal standard. After extraction, samples were centrifuged (2500 x g, 10 min) and the supernatant evaporated to dryness under a nitrogen stream. Extracted lipids were reconstituted in 400 µl tetrahydrofuran:methanol:water (7:2:1, v/v/v). UPLC-nanoESI-MS/MS analysis was carried out as described (Herrfurth et al., 2021) with the parameters listed in Suppl. Table S2. Peak integration was performed with the MultiQuant software (AB Sciex, Framingham, MA, USA). Quantitative analysis of integrated peak values was done in RStudio 4.0.1.

For the analysis of the free sterols, the samples were extracted in the same manner and heptadecanoic acid was added as an internal standard. Aliquots of the samples (1/8 of the volume) were dried under nitrogen stream, dissolved in 15 µl pyridine and derivatized with 30 µl N-methyl-N-(trimethylsilyl)trifluoracetamid (MSTFA) before being analyzed by GC-MS (Agilent 7890B GC-Agilent 5977N-MSD) as previously described (Berghoff et al., 2021).

Phospho- and galactolipids were also extracted from frozen and homogenized leaf material. To inactivate phospholipase activity, samples were initially incubated in boiling propan-2-ol for 20 min. After evaporating the leaf material to dryness, lipids were sequentially extracted with 1 ml of chloroform:methanol (1:2, v/v), 1 ml of chloroform:methanol (2:1, v/v), and 1 ml of chloroform. For each extraction step, samples were vortexed thoroughly, centrifuged for 10 min at 1500 x g and the supernatants collected in a new tube. To the combined supernatant, 0.75 ml of 300 mM NH_4_CH_3_CO_2_ was added, samples were vortexed thoroughly, centrifuged for 5 min at 1500 x g, and the lower phase transferred to a new tube. Extracts were evaporated to dryness and dissolved in chloroform:methanol:300 mM ammonium acetate (300:665:35, v/v/v). The dry weight of the sample was determined and the amount of the internal standard adjusted accordingly. Samples were analyzed via direct infusion nanospray MS on an Agilent 6530 Accurate-Mass Q-TOF LC/MS instrument equipped with a ChipCube interface as previously described (Welti et al., 2002; Gasulla et al., 2013; Gutbrod et al., 2021). The parameters for the MS/MS experiments are listed in Suppl. Dataset S1.

### Proteomic sample preparation and LC-MS/MS analysis of peptides

Proteins were defatted and washed with 80% ethanol and then dissolved in 6 M urea and 5% (w/v) SDS. Determination of protein concentration, in-gel tryptic-digest and peptide desalting was carried out as described previously (Shevchenko et al., 2006; Rappsilber et al., 2007; Kretzschmar et al., 2018). Dried peptides were reconstituted in 20 µl sample buffer (2% acetonitrile, 0.1% formic acid) and subjected to LC-MS/MS analysis. To that end, 1 to 3 µl of each sample were subjected to reverse phase LC for peptide separation using an RSLCnano Ultimate 3000 system (Thermo Fisher Scientific). Peptides were loaded on an Acclaim PepMap 100 pre-column (100 µm x 2 cm, C18, 5 µm, 100 Å; Thermo Fisher Scientific) with 0.07% trifluoroacetic acid at a flow rate of 20 µL/min for 3 min. Analytical separation of peptides was accomplished on an Acclaim PepMap RSLC column (75 µm x 50 cm, C18, 2 µm, 100 Å; Thermo Fisher Scientific) at a flow rate of 300 nL/min. The solvent composition followed a gradual change within 94 min: from 96% solvent A (0.1% formic acid) and 4% solvent B (80% acetonitrile, 0.1% formic acid) to 10% solvent B within 2 minutes, to 30% solvent B within the next 58 min, to 45% solvent B within the following 22 min, and to 90% solvent B within the last 12 min of the gradient. All solvents and acids had Optima grade for LC-MS (Thermo Fisher Scientific). Eluting peptides were on-line ionized by nano-electrospray (nESI) using the Nanospray Flex Ion Source (Thermo Fisher Scientific) at 1.5 kV (liquid junction) and transferred to the mass spectrometer. For mass spectrometry, either an Orbitrap Velos Pro hybrid mass spectrometer or a Q Exactive HF mass spectrometer (both Thermo Fisher Scientific) were used. On the Orbitrap Velos Pro hybrid mass spectrometer, full scans were recorded within a mass range of 300 to 1850 m/z at a resolution of 30,000 with the Orbitrap-FT analyzer. Full scans were followed by data-dependent top 10 CID fragmentation (dynamic exclusion enabled) within the ion trap Velos Pro analyzer. For the Q Exactive HF mass spectrometer, full scans were recorded in a mass range of 300 to 1650 m/z at a resolution of 30,000 and followed by data-dependent top 10 HCD fragmentation at a resolution of 15,000 (dynamic exclusion enabled). LC-MS method programming and data acquisition was performed with the XCalibur 4.0 software (Thermo Fisher Scientific).

### Computational processing of MS/MS data

MS/MS raw data were processed in the MaxQuant software (version 1.6.2.17) for feature detection, peptide identification and protein group assembly (Cox and Mann, 2008). Mostly, default settings were used with additional settings as specified in Suppl. Table S1. The TAIR10 protein database was used as reference for identification. For quantification, label free quantification was calculated according to the iBAQ and LFQ algorithms (Cox and Mann, 2008; Schwanhäusser et al., 2011; Cox et al., 2014). Further data analysis was done in Perseus 1.6.2.2 (Tyanova et al., 2016), Excel (Microsoft Office) and RStudio 4.0.1 (RStudio Team (2021). RStudio: Integrated Development Environment for R. RStudio, PBC, Boston, MA. http://www.rstudio.com/). All relevant data can be found within the manuscript and its supporting materials. Proteomic raw data can be found in the PRIDE database (Vizcaíno et al., 2014) under the identifier PXD045596 (https://www.ebi.ac.uk/pride/). All metadata can be found in Suppl. Table S1.

Protein localization was annotated based on the Plant Proteome Database (Sun et al., 2009) as of 14^th^ June 2022. LD localization was assigned based on previous studies (Kretzschmar et al., 2018; Fernández-Santos et al., 2020; Kretzschmar et al., 2020; Doner et al., 2021; Ge et al., 2022; Li et al., 2022). Volcano plots were calculated in Perseus. rLFQ and riBAQ values of proteins were log2-transformed and missing values were imputed from normal distribution (parameters: width 0.3, down shift 2.5, separately for each sample). For the comparison of different treatments, plant lines or isolated fraction, *p*-values were calculated by two-sided t-tests. For plotting, transformed and imputed rLFQ or riBAQ values, and calculated *p*-values were exported and plotted in RStudio 4.0.1.

Proteins with relevance in lipid metabolism were selected based on a compiled list of Arabidopsis lipid genes using KEGG pathway (https://www.genome.jp/kegg/pathway.html, latest update 10th March 2020), Aralip (http://aralip.plantbiology.msu.edu/pathways/pathways; (Li-Beisson et al., 2013) and genes from (Kelly and Feussner, 2016); (Kretzschmar et al., 2020); and from Plant Sphingolipid Metabolism and Function, Chapter 11 (Luttgeharm et al., 2016). n.d., not detected in control and treatment, n.d. in control, only detected under treatment. p-values are based on Student’s t-test.

Identification of candidate proteins for LD localization was done using web tools. Potential membrane-spanning regions were predicted using the TMHMM - 2.0 server, which predicts transmembrane helices based on hidden Markov models (Krogh et al., 2001).

### PR gene expression analysis via qRT-PCR

Leaves of wild-type and *tgd1-1 sdp1-4* plants were frozen in liquid nitrogen and ground to fine powder with a ball mill (Retsch GmbH, Haan, Germany). RNA was extracted from 100 mg of ground leaf material per biological replicate and line. RNA isolation was done using the Spectrum Plant Total RNA Kit (Sigma-Aldrich) and subsequently treated with DNaseI (Thermo Fisher Scientific, Waltham, MA, USA) according to manufacturer’s instructions. For cDNA synthesis, 0.5 µg RNA was reverse transcribed (Maxima Reverse Transcriptase; Thermo Fisher Scientific). The reaction product was diluted 1:10 in double-distilled water before qPCR. For each qPCR sample, 4 µl of the diluted cDNA was used together with Takyon No Rox SYBR MasterMix dTTP Blue (Eurogentec, Seraing, Belgium). *AT3G01150* was chosen as reference gene (Czechowski et al., 2005), used primers for all genes of interest and the reference gene are listed in Suppl. Table S3. The following PCR program was used for amplification: 95°C for 1 min 20 s (95°C for 20 s, 58°C for 20 s, 72°C for 40 s) × 39, 72°C 4 min. Amplicons were tested by melt curve analysis. PCR amplification and melt curve analysis were carried out in an iQ5 qPCR cycler (BioRad Laboratories, Hercules, CA, USA).

### Analysis of phytohormone levels

Phytohormones were extracted by biphasic extraction with methyl-*tert*-butyl ether based on (Matyash et al., 2008). After extraction, phytohormones were reversed phase-separated using an ACQUITY UPLC system (Waters Corp., Milford, MA, USA) and analyzed by nanoelectrospray ionization (nanoESI) (TriVersa Nanomate, Advion BioSciences, Ithaca, NY, USA) coupled with an AB Sciex 4000 QTRAP tandem mass spectrometer (AB Sciex, Framingham, MA, USA) employed in scheduled multiple reaction monitoring modes (Herrfurth and Feussner, 2020) with the following modifications. For quantification, 10 ng D_4_-SA (C/D/N Isotopes Inc., Pointe-Claire, Canada) were added at the beginning of the extraction procedure. For SA and SAG analysis, the following mass transitions were included: 137/93 [declustering potential (DP) -25 V, entrance potential (EP) -6 V, collision energy (CE) -20 V] for SA, 141/97 (DP -25 V, EP -6 V, CE -22 V) for D_4_-SA, and 299/137 (DP -30 V, EP -4 V, CE -18 V) for SAG.

### Molecular cloning and candidate localization studies in Nicotiana benthamiana leaves

Open reading frames of selected candidate genes were amplified from cDNA prepared with Maxima Reverse Transcriptase (Thermo Fisher Scientific) according to manufactureres instruction using leaf RNA that had been extracted using the Spectrum Plant Total RNA Kit (Merck KGaA, Darmstadt, Germany). Constructs were amplified with the Phusion High-Fidelity DNA Polymerase (Thermo Fisher Scientific, Waltham, MA, USA) following manufacturer’s instructions and using primers listed in Suppl. Table S4. Gateway cloning into the plant binary vectors pMDC32-ChC and pMDC32-ChN was carried out by traditional or fast Gateway® cloning as described in (Müller et al., 2017). Vector construction of pMDC32-ChC and pMDC32-ChN has been described previously in Kretzschmar et al., 2020 and Doner et al., 2021, respectively. The construct of MmDGAT2 in pMDC32, which was used for co-expression experiments, has been described in Cai et al., 2019. Localization of candidates was analyzed in leaves of *N. benthamiana* that were transiently transformed by infiltration with *Agrobacterium tumefaciens* harboring candidate expression vectors. *N. benthamiana* plant growth, leaf infiltration, BODIPY staining, and CLSM imaging was performed as previously described (Gidda et al., 2016; Kretzschmar et al., 2020). Micrographs of leaves were acquired using a Leica SP5 CLSM (Leica Microsystems). Excitations and emission signals for fluorescent proteins and BODIPY were collected sequentially as single optical sections in double-labeling experiments like those described in Gidda et al., 2016.

## Accession numbers

AT3G01420 (α-DOX1); AT1G73680 (α-DOX2); AT2G07050 (CAS); AT5G53560 (CB5-E); AT2G33380 (CLO3); AT2G17840 (ERD7); AT1G01610 (GPAT4); AT4G00400 (GPAT8); AT1G67360 (LDAP1); AT2G47780 (LDAP2); AT3G05500 (LDAP3); AT5G16550 (LDIP); AT5G04830 (LDNP); AT1G43890 (LDS1); AT1G73920 (LIDL2); AT4G33110 (LIME1); AT4G33120 (LIME2); AT1G45201 (OBL3); AT4G10790 (PUX10); AT5G13710 (SMT1); NP_080660.1 (MmDGAT2)

## Supplemental Data

**Supplemental Figure S1:** Membrane and sterol lipid composition in heat-stressed leaves.

**Supplemental Figure S2:** Membrane lipid composition after infection by *B. cinerea*.

**Supplemental Figure S3:** Composition of membrane lipids after infection with *P. syringae*. **Supplemental Figure S4:** Relative composition of triacylglycerols and sterol esters species after different infection treatments.

**Supplemental Figure S5:** STRING networks of differentially accumulating proteins of Arabidopsis Col-0 leaves after different stresses.

**Supplemental Figure S6:** Alterations in total abundance and species composition of membrane and hydrophobic lipids of the Arabidopsis double mutant *tgd1-1 sdp1-4*.

**Supplemental Figure S7:** Alterations in species composition of membrane and sterol lipids of the Arabidopsis double mutant *tgd1-1 sdp1-4*.

**Supplemental Figure S8:** STRING networks of differentially accumulating proteins in leaf total protein fractions of Arabidopsis Col-0 and *tgd1-1 sdp1-4*.

**Supplemental Figure S9:** Protein abundance of three pathogenesis-related proteins in Arabidopsis leaves of Col-0 *and tgd1-1 sdp1-4* after different treatments.

**Supplemental Figure S10:** Enrichment of different organelle proteomes in the LD-enriched fraction.

**Supplemental Figure S11:** Prediction of the membrane-interacting region in CB5-E.

**Supplemental Figure S12:** Additional subcellular localization studies of candidate proteins in *Nicotiana benthamiana* leaves.

**Supplemental Table S1:** Metadata file for LC-MS/MS data processing with MaxQuant.

**Supplemental Table S2:** Parameters for lipid analysis by UPLC-nanoESI-MS/MS.

**Supplemental Table S3:** Primers used for gene expression analysis via qPCR

**Supplemental Table S4:** Primers used for Gateway cloning and sequencing

**Supplemental Dataset S1:** Arabidopsis leaf membrane lipids - absolute amounts.

**Supplemental Dataset S2:** Arabidopsis leaf membrane lipids - relative composition of species per class.

**Supplemental Dataset S3:** Arabidopsis leaf neutral lipids - absolute peak areas.

**Supplemental Dataset S4:** Arabidopsis leaf neutral lipids - normalized icf-corrected peak areas.

**Supplemental Dataset S5:** Relative contribution of sterol esters with a common sterol moiety to total sterol ester signal intensity.

**Supplemental Dataset S6:** Relative proportions of individual TAG species.

**Supplemental Dataset S7:** Free sterols

**Supplemental Dataset S8:** Proteins found in Arabidopsis leaves - normalised rLFQ and riBAQ values.

**Supplemental Dataset S9:** Comparison of protein abundance in Arabidopsis Col-0 leaves after infection with *B. cinerea* to mock-treated plants.

**Supplemental Dataset S10:** Comparison of protein abundance in Arabidopsis Col-0 leaves in reaction to infection with *P. syringae* pv. *tomato* DC3000 *ΔavrPto ΔavrPtoB*.

**Supplemental Dataset S11:** Comparison of protein abundance in Arabidopsis Col-0 leaves after heat stress.

**Supplemental Dataset S12:** Proteins with relevance in lipid metabolism found in Arabidopsis leaves - normalised rLFQ and riBAQ values

**Supplemental Dataset S13:** Comparison of protein abundance in Arabidopsis *tgd1-1 sdp1-4* leaves after infection with *B. cinerea* to mock-treated plants.

**Supplemental Dataset S14:** Comparison of protein abundance in Arabidopsis *tgd1-1 sdp1-4* leaves in reaction to infection with *P. syringae* pv. *tomato* DC3000 *ΔavrPto ΔavrPtoB*.

**Supplemental Dataset S15:** Comparison of protein abundance in Arabidopsis *tgd1-1 sdp1-4* leaves after heat stress.

**Supplemental Dataset S16:** Comparison of proteins in non-stressed leaves of Arabidopsis Col-0 and *tgd1-1 sdp1-4*.

**Supplemental Dataset S17:** Comparison of proteins in LD-enriched fractions to total protein fractions of Arabidopsis *tgd1-1 sdp1-4* leaves.

## Acknowledgements

We are grateful to Marcel Wiermer for his support with pathogen infections and we thank him and George Haughn for valuable advice. We thank Changcheng Xu for providing the *tgd1-1 sdp1-4* seeds. Technical assistance was provided by Helga Peisker, Jannis Anstatt, Denis Krone, and Annabel Maisl.

## List of author contributions

P.S., J.S., R.T.M. and T.I. designed the work, P.S., N.M.D., K.G., C.H., P.N., M.S.S.L., K.B., K.S. and O.V. performed research, P.S., N.M.D., K.G., C.H., K.S., I.F., P.D., G.H.B., R.T.M. and T.I. analyzed data, and P.S., R.T.M. and T.I. wrote the manuscript with contributions by the other authors. All authors critically read and revised the manuscript and approved the final version.

## References

Ahmad L, Rylott EL, Bruce NC, Edwards R, Grogan G (2017) Structural evidence for Arabidopsis glutathione transferase AtGSTF2 functioning as a transporter of small organic ligands. FEBS Open Bio 7: 122–132

Aubert Y, Vile D, Pervent M, Aldon D, Ranty B, Simonneau T, Vavasseur A, Galaud J-P (2010) RD20, a stress-inducible caleosin, participates in stomatal control, transpiration and drought tolerance in Arabidopsis thaliana. Plant Cell Physiol 51: 1975–1987

Bauwe H, Hagemann M, Fernie AR (2010) Photorespiration: players, partners and origin. Trends Plant Sci 15: 330–336

Berardini TZ, Reiser L, Li D, Mezheritsky Y, Muller R, Strait E, Huala E (2015) The Arabidopsis information resource: Making and mining the “gold standard” annotated reference plant genome. Genesis 53: 474–485

Berghoff SA, Spieth L, Sun T, Hosang L, Depp C, Sasmita AO, Vasileva MH, Scholz P, Zhao Y, Krueger-Burg D, et al (2021) Neuronal cholesterol synthesis is essential for repair of chronically demyelinated lesions in mice. Cell Rep 37: 109889

Bilgin DD, Zavala JA, Zhu J, Clough SJ, Ort DR, DeLucia EH (2010) Biotic stress globally downregulates photosynthesis genes. Plant Cell Environ 33: 1597–1613

Blée E, Boachon B, Burcklen M, Le Guédard M, Hanano A, Heintz D, Ehlting J, Herrfurth C, Feussner I, Bessoule J-J (2014) The reductase activity of the Arabidopsis caleosin RESPONSIVE TO DESSICATION20 mediates gibberellin-dependent flowering time, abscisic acid sensitivity, and tolerance to oxidative stress. Plant Physiol 166: 109–124

Bonfig KB, Schreiber U, Gabler A, Roitsch T, Berger S (2006) Infection with virulent and avirulent P. syringae strains differentially affects photosynthesis and sink metabolism in Arabidopsis leaves. Planta 225: 1–12

Böttcher C, Westphal L, Schmotz C, Prade E, Scheel D, Glawischnig E (2009) The multifunctional enzyme CYP71B15 (PHYTOALEXIN DEFICIENT3) converts cysteine-indole-3-acetonitrile to camalexin in the indole-3-acetonitrile metabolic network of *Arabidopsis thaliana*. Plant Cell 21: 1830–1845

Brocard L, Immel F, Coulon D, Esnay N, Tuphile K, Pascal S, Claverol S, Fouillen L, Bessoule J-J, Bréhélin C (2017) Proteomic analysis of lipid droplets from Arabidopsis aging leaves brings new insight into their biogenesis and functions. Front Plant Sci 8: 894

Cai Y, Goodman JM, Pyc M, Mullen RT, Dyer JM, Chapman KD (2015) Arabidopsis SEIPIN proteins modulate triacylglycerol accumulation and influence lipid droplet proliferation. Plant Cell 27: 2616–2636

Cai Y, Whitehead P, Chappell J, Chapman KD (2019) Mouse lipogenic proteins promote the co-accumulation of triacylglycerols and sesquiterpenes in plant cells. Planta 250: 79–94

Cernac A, Benning C (2004) WRINKLED1 encodes an AP2/EREB domain protein involved in the control of storage compound biosynthesis in Arabidopsis. Plant J 40: 575–585

Chen JCF, Tsai CCY, Tzen JTC (1999) Cloning and secondary structure analysis of caleosin, a unique calcium-binding protein in oil bodies of plant seeds. Plant Cell Physiol 40: 1079–1086

Corey EJ, Matsuda SP, Bartel B (1993) Isolation of an Arabidopsis thaliana gene encoding cycloartenol synthase by functional expression in a yeast mutant lacking lanosterol synthase by the use of a chromatographic screen. Proc Natl Acad Sci U S A 90: 11628–11632

Cox J, Hein MY, Luber CA, Paron I, Nagaraj N, Mann M (2014) Accurate proteome-wide label-free quantification by delayed normalization and maximal peptide ratio extraction, termed MaxLFQ. Mol Cell Proteomics 13: 2513–2526

Cox J, Mann M (2008) MaxQuant enables high peptide identification rates, individualized p.p.b.-range mass accuracies and proteome-wide protein quantification. Nat Biotechnol 26: 1367–1372

Crawford T, Lehotai N, Strand Å (2018) The role of retrograde signals during plant stress responses. J Exp Bot 69: 2783–2795

Cummins I, Hills MJ, Ross JHE, Hobbs DH, Watson MD, Murphy DJ (1993) Differential, temporal and spatial expression of genes involved in storage oil and oleosin accumulation in developing rapeseed embryos: implications for the role of oleosins and the mechanisms of oil-body formation. Plant Mol Biol 23: 1015–1027

Czechowski T, Stitt M, Altmann T, Udvardi MK, Scheible W-R (2005) Genome-wide identification and testing of superior reference genes for transcript normalization in Arabidopsis. Plant Physiol 139: 5–17

Deruyffelaere C, Purkrtova Z, Bouchez I, Collet B, Cacas J-L, Chardot T, Gallois J-L, D’Andrea S (2018) PUX10 is a CDC48A adaptor protein that regulates the extraction of ubiquitinated oleosins from seed lipid droplets in Arabidopsis. Plant Cell 30: 2116–2136

Diener AC, Li H, Zhou W, Whoriskey WJ, Nes WD, Fink GR (2000) Sterol methyltransferase 1 controls the level of cholesterol in plants. Plant Cell 12: 853–870

Dixon DP, Skipsey M, Edwards R (2010) Roles for glutathione transferases in plant secondary metabolism. Phytochemistry 71: 338–350

Doner NM, Seay D, Mehling M, Sun S, Gidda SK, Schmitt K, Braus GH, Ischebeck T, Chapman KD, Dyer JM, et al (2021) Arabidopsis thaliana EARLY RESPONSIVE TO DEHYDRATION 7 localizes to lipid droplets via its senescence domain. Front Plant Sci 12: 658961

Dufourc EJ (2008) Sterols and membrane dynamics. J Chem Biol 1: 63–77

Eberhardt RY, Chang Y, Bateman A, Murzin AG, Axelrod HL, Hwang WC, Aravind L (2013) Filling out the structural map of the NTF2-like superfamily. BMC Bioinformatics 14: 327

Falcone DL, Ogas JP, Somerville CR (2004) Regulation of membrane fatty acid composition by temperature in mutants of Arabidopsis with alterations in membrane lipid composition. BMC Plant Biol 4: 17

Fan J, Yan C, Roston R, Shanklin J, Xu C (2014) Arabidopsis lipins, PDAT1 acyltransferase, and SDP1 triacylglycerol lipase synergistically direct fatty acids toward β-oxidation, thereby maintaining membrane lipid homeostasis. Plant Cell 26: 4119–4134

Fernández-Santos R, Izquierdo Y, López A, Muñiz L, Martínez M, Cascón T, Hamberg M, Castresana C (2020) Protein profiles of lipid droplets during the hypersensitive defense response of Arabidopsis against Pseudomonas infection. Plant Cell Physiol 61: 1144–1157

Fu ZQ, Dong X (2013) Systemic acquired resistance: Turning local infection into global defense. Annu Rev Plant Biol 64: 839–863

Fukuchi-Mizutani M, Mizutani M, Tanaka Y, Kusumi T, Ohta D (1999) Microsomal electron transfer in higher plants: Cloning and heterologous expression of NADH-cytochrome *b* 5 reductase from Arabidopsis. Plant Physiol 119: 353–362

Gasulla F, Vom Dorp K, Dombrink I, Zähringer U, Gisch N, Dörmann P, Bartels D (2013) The role of lipid metabolism in the acquisition of desiccation tolerance in Craterostigma plantagineum: a comparative approach. Plant J 75: 726–741

Ge S, Zhang R-X, Wang Y-F, Sun P, Chu J, Li J, Sun P, Wang J, Hetherington AM, Liang Y-K (2022) The Arabidopsis Rab protein RABC1 affects stomatal development by regulating lipid droplet dynamics. Plant Cell 34: 4274–4292

Germain V, Rylott EL, Larson TR, Sherson SM, Bechtold N, Carde J-P, Bryce JH, Graham IA, Smith SM (2001) Requirement for 3-ketoacyl-CoA thiolase-2 in peroxisome development, fatty acid β-oxidation and breakdown of triacylglycerol in lipid bodies of Arabidopsis seedlings. Plant J 28: 1–12

Gidda SK, Park S, Pyc M, Yurchenko O, Cai Y, Wu P, Andrews DW, Chapman KD, Dyer JM, Mullen RT (2016) Lipid droplet-associated proteins (LDAPs) are required for the dynamic regulation of neutral lipid compartmentation in plant cells. Plant Physiol 170: 2052–2071

Griebel T, Zeier J (2010) A role for beta-sitosterol to stigmasterol conversion in plant-pathogen interactions. Plant J 63: 254–268

Gutbrod K, Peisker H, Dörmann P (2021) Direct infusion mass spectrometry for complex lipid analysis. In D Bartels, P Dörmann, eds, Plant Lipids. Springer US, New York, NY, pp 101–115

Guzha A, McGee R, Scholz P, Hartken D, Lüdke D, Bauer K, Wenig M, Zienkiewicz K, Herrfurth C, Feussner I, et al (2022) Cell wall-localized BETA-XYLOSIDASE4 contributes to immunity of Arabidopsis against *Botrytis cinerea*. Plant Physiol 189: 1794–1813

Guzha A, Whitehead P, Ischebeck T, Chapman KD (2023) Lipid droplets: Packing hydrophobic molecules within the aqueous cytoplasm. Annu Rev Plant Biol 74: 195–223

Hemmingsen SM, Woolford C, van der Vies SM, Tilly K, Dennis DT, Georgopoulos CP, Hendrix RW, Ellis RJ (1988) Homologous plant and bacterial proteins chaperone oligomeric protein assembly. Nature 333: 330–334

Herrfurth C, Feussner I (2020) Quantitative jasmonate profiling using a high-throughput UPLC-NanoESI-MS/MS method. Methods Mol Biol 2085: 169–187

Herrfurth C, Liu Y-T, Feussner I (2021) Targeted analysis of the plant lipidome by UPLC-NanoESI-MS/MS. Methods Mol Biol 2295: 135–155

Higashi Y, Okazaki Y, Myouga F, Shinozaki K, Saito K (2015) Landscape of the lipidome and transcriptome under heat stress in Arabidopsis thaliana. Sci Rep 5: 1–11

Higashi Y, Saito K (2019) Lipidomic studies of membrane glycerolipids in plant leaves under heat stress. Prog Lipid Res 75: 100990

Hsieh K, Huang AHC (2004) Endoplasmic reticulum, oleosins, and oils in seeds and tapetum cells. Plant Physiol 136: 3427–3434

Ischebeck T, Krawczyk HE, Mullen RT, Dyer JM, Chapman KD (2020) Lipid droplets in plants and algae: Distribution, formation, turnover and function. Semin Cell Dev Biol 108: 82–93

Ishikawa A, Tanaka H, Nakai M, Asahi T (2003) Deletion of a chaperonin 60β gene leads to cell death in the Arabidopsis lesion initiation 1 mutant. Plant Cell Physiol 44: 255–261

Jiang Z, He F, Zhang Z (2017) Large-scale transcriptome analysis reveals arabidopsis metabolic pathways are frequently influenced by different pathogens. Plant Mol Biol 94: 453–467

Jolivet P, Boulard C, Bellamy A, Larré C, Barre M, Rogniaux H, d’Andréa S, Chardot T, Nesi N (2009) Protein composition of oil bodies from mature Brassica napus seeds. Proteomics 9: 3268–3284

Jolivet P, Roux E, D’Andrea S, Davanture M, Negroni L, Zivy M, Chardot T (2004) Protein composition of oil bodies in Arabidopsis thaliana ecotype WS. Plant Physiol Biochem 42: 501– 509

Joyard J, Maréchal E, Miège C, Block MA, Dorne A-J, Douce R (2004) Structure, distribution and biosynthesis of glycerolipids from higher plant chloroplasts. *In* P-A Siegenthaler, M Norio, eds, Lipids in Photosynthesis: Structure, Function and Genetics. Kluwer Academic Publishers, Dordrecht, pp 21–52

Katavic V, Agrawal GK, Hajduch M, Harris SL, Thelen JJ (2006) Protein and lipid composition analysis of oil bodies from two Brassica napus cultivars. Proteomics 6: 4586–4598

Kelly AA, Feussner I (2016) Oil is on the agenda: Lipid turnover in higher plants. Biochim Biophys Acta 1861: 1253–1268

Khor VK, Ahrends R, Lin Y, Shen W-J, Adams CM, Roseman AN, Cortez Y, Teruel MN, Azhar S, Kraemer FB (2014) The proteome of cholesteryl-ester-enriched versus triacylglycerol-enriched lipid droplets. PLoS One 9: e105047

Kim EY, Park KY, Seo YS, Kim WT (2016a) Arabidopsis Small Rubber Particle Protein Homolog SRPs Play Dual Roles as Positive Factors for Tissue Growth and Development and in Drought Stress Responses. Plant Physiol 170: 2494–2510

Kim EY, Park KY, Seo YS, Kim WT (2016b) Arabidopsis small rubber particle protein homolog SRPs play dual roles as positive factors for tissue growth and development and in drought stress responses. Plant Physiol 170: 2494–2510

Kim HU, Lee K-R, Jung S-J, Shin HA, Go YS, Suh M-C, Kim JB (2015) Senescence-inducible LEC2 enhances triacylglycerol accumulation in leaves without negatively affecting plant growth. Plant Biotechnol J 13: 1346–1359

Kim K, Portis AR (2005) Temperature dependence of photosynthesis in Arabidopsis plants with modifications in Rubisco Activase and membrane fluidity. Plant Cell Physiol 46: 522–530

Klepikova AV, Kasianov AS, Gerasimov ES, Logacheva MD, Penin AA (2016) A high resolution map of the Arabidopsis thaliana developmental transcriptome based on RNA-seq profiling. Plant J 88: 1058–1070

Kliebenstein DJ, Rowe HC, Denby KJ (2005) Secondary metabolites influence Arabidopsis/Botrytis interactions: variation in host production and pathogen sensitivity. Plant J 44: 25–36

Kobza J, Edwards GE (1987) Influences of leaf temperature on photosynthetic carbon metabolism in wheat. Plant Physiol 83: 69–74

Kovacs D, Kalmar E, Torok Z, Tompa P (2008) Chaperone activity of ERD10 and ERD14, two disordered stress-related plant proteins. Plant Physiol 147: 381–390

Krawczyk HE, Rotsch AH, Herrfurth C, Scholz P, Shomroni O, Salinas-Riester G, Feussner I, Ischebeck T (2022a) Heat stress leads to rapid lipid remodeling and transcriptional adaptations in Nicotiana tabacum pollen tubes. Plant Physiol 189: 490–515

Krawczyk HE, Sun S, Doner NM, Yan Q, Lim MSS, Scholz P, Niemeyer PW, Schmitt K, Valerius O, Pleskot R, et al (2022b) SEED LIPID DROPLET PROTEIN1, SEED LIPID DROPLET PROTEIN2, and LIPID DROPLET PLASMA MEMBRANE ADAPTOR mediate lipid droplet-plasma membrane tethering. Plant Cell 34: 2424–2448

Kretzschmar FK, Doner NM, Krawczyk HE, Scholz P, Schmitt K, Valerius O, Braus GH, Mullen RT, Ischebeck T (2020) Identification of low-abundance lipid droplet proteins in seeds and seedlings. Plant Physiol 182: 1326–1345

Kretzschmar FK, Mengel LA, Müller AO, Schmitt K, Blersch KF, Valerius O, Braus GH, Ischebeck T (2018) PUX10 is a lipid droplet-localized scaffold protein that interacts with CELL DIVISION CYCLE48 and is involved in the degradation of lipid droplet proteins. Plant Cell 30: 2137–2160

Krogh A, Larsson B, von Heijne G, Sonnhammer EL (2001) Predicting transmembrane protein topology with a hidden Markov model: application to complete genomes. J Mol Biol 305: 567– 580

Kumar D, Hazra S, Datta R, Chattopadhyay S (2016) Transcriptome analysis of Arabidopsis mutants suggests a crosstalk between ABA, ethylene and GSH against combined cold and osmotic stress. Sci Rep 6: 36867

Kumar R, Tran L-SP, Neelakandan AK, Nguyen HT (2012) Higher plant cytochrome b5 polypeptides modulate fatty acid desaturation. PLoS One 7: e31370

Kumar R, Wallis JG, Skidmore C, Browse J (2006) A mutation in Arabidopsis cytochrome b5 reductase identified by high-throughput screening differentially affects hydroxylation and desaturation. Plant J 48: 920–932

La Camera S, Geoffroy P, Samaha H, Ndiaye A, Rahim G, Legrand M, Heitz T (2005) A pathogen-inducible patatin-like lipid acyl hydrolase facilitates fungal and bacterial host colonization in Arabidopsis. Plant J 44: 810–825

Li F, Han X, Guan H, Xu MC, Dong YX, Gao X-Q (2022) PALD encoding a lipid droplet-associated protein is critical for the accumulation of lipid droplets and pollen longevity in Arabidopsis. New Phytol 235: 204–219

Li-Beisson Y, Shorrosh B, Beisson F, Andersson MX, Arondel V, Bates PD, Baud S, Bird D, Debono A, Durrett TP, et al (2013) Acyl-lipid metabolism. Arabidopsis Book 11: e0161

Lin BL, Wang JS, Liu HC, Chen RW, Meyer Y, Barakat A, Delseny M (2001) Genomic analysis of the Hsp70 superfamily in Arabidopsis thaliana. Cell Stress Chaperones 6: 201–208

Lin L-J, Tai SSK, Peng C-C, Tzen JTC (2002) Steroleosin, a sterol-binding dehydrogenase in seed oil bodies. Plant Physiol 128: 1200–1211

Lin Y-T, Chen L-J, Herrfurth C, Feussner I, Li H (2016) Reduced biosynthesis of digalactosyldiacylglycerol, a major chloroplast membrane lipid, leads to oxylipin overproduction and phloem cap lignification in Arabidopsis. Plant Cell 28: 219–232

Listenberger LL, Brown DA (2007) Fluorescent detection of lipid droplets and associated proteins. Curr Protoc Cell Biol 35: 24.2.1–24.2.11

Luttgeharm KD, Kimberlin AN, Cahoon EB (2016) Plant sphingolipid metabolism and function. *In* Y Nakamura, Y Li-Beisson, eds, Lipids in Plant and Algae Development. Springer International Publishing, Cham, pp 249–286

Maggio C, Barbante A, Ferro F, Frigerio L, Pedrazzini E (2007) Intracellular sorting of the tail-anchored protein cytochrome b5 in plants: a comparative study using different isoforms from rabbit and Arabidopsis. J Exp Bot 58: 1365–1379

Matyash V, Liebisch G, Kurzchalia TV, Shevchenko A, Schwudke D (2008) Lipid extraction by methyl-*tert*-butyl ether for high-throughput lipidomics. J Lipid Res 49: 1137–1146

Moellering ER, Muthan B, Benning C (2010) Freezing tolerance in plants requires lipid remodeling at the outer chloroplast membrane. Science 330: 226–228

Morikawa T, Mizutani M, Aoki N, Watanabe B, Saga H, Saito S, Oikawa A, Suzuki H, Sakurai N, Shibata D, et al (2006) Cytochrome P450 CYP710A encodes the sterol C-22 desaturase in Arabidopsis and tomato. Plant Cell 18: 1008–1022

Mueller SP, Krause DM, Mueller MJ, Fekete A (2015) Accumulation of extra-chloroplastic triacylglycerols in Arabidopsis seedlings during heat acclimation. J Exp Bot 66: 4517–4526

Mueller SP, Unger M, Guender L, Fekete A, Mueller MJ (2017) Phospholipid:diacylglycerol acyltransferase-mediated triacylglyerol synthesis augments basal thermotolerance. Plant Physiol 175: 486–497

Mukherjee AK, Carp M-J, Zuchman R, Ziv T, Horwitz BA, Gepstein S (2010) Proteomics of the response of Arabidopsis thaliana to infection with Alternaria brassicicola. J Proteomics 73: 709–720

Müller AO, Blersch KF, Gippert AL, Ischebeck T (2017) Tobacco pollen tubes – a fast and easy tool for studying lipid droplet association of plant proteins. Plant J 89: 1055–1064

Müller AO, Ischebeck T (2018) Characterization of the enzymatic activity and physiological function of the lipid droplet-associated triacylglycerol lipase AtOBL1. New Phytol 217: 1062–1076

Murashige T, Skoog F (1962) A revised medium for rapid growth and bio assays with tobacco tissue cultures. Physiol Plant 15: 473–497

Murphy DJ (1993) Structure, function and biogenesis of storage lipid bodies and oleosins in plants. Prog Lipid Res 32: 247–280

Partridge M, Murphy DJ (2009) Roles of a membrane-bound caleosin and putative peroxygenase in biotic and abiotic stress responses in Arabidopsis. Plant Physiol Biochem 47: 796–806

Pavicic M, Overmyer K, Rehman A ur, Jones P, Jacobson D, Himanen K (2021) Image-based methods to score fungal pathogen symptom progression and severity in excised Arabidopsis leaves. Plants (Basel) 10: 158

Pieterse CMJ, Van der Does D, Zamioudis C, Leon-Reyes A, Van Wees SCM (2012) Hormonal modulation of plant immunity. Annu Rev Cell Dev Biol 28: 489–521

Piotrowski M, Schönfelder S, Weiler EW (2001) The Arabidopsis thaliana isogene NIT4 and its orthologs in tobacco encode beta-cyano-L-alanine hydratase/nitrilase. J Biol Chem 276: 2616–2621

Pyc M, Cai Y, Gidda SK, Yurchenko O, Park S, Kretzschmar FK, Ischebeck T, Valerius O, Braus GH, Chapman KD, et al (2017) Arabidopsis lipid droplet-associated protein (LDAP) - interacting protein (LDIP) influences lipid droplet size and neutral lipid homeostasis in both leaves and seeds. Plant J 92: 1182–1201

Pyc M, Gidda SK, Seay D, Esnay N, Kretzschmar FK, Cai Y, Doner NM, Greer MS, Hull JJ, Coulon D, et al (2021) LDIP cooperates with SEIPIN and LDAP to facilitate lipid droplet biogenesis in Arabidopsis. Plant Cell 33: 3076–3103

Qi Y, Katagiri F (2012) Membrane microdomain may be a platform for immune signaling. Plant Signal Behav 7: 454–456

Qi Y, Tsuda K, Nguyen LV, Wang X, Lin J, Murphy AS, Glazebrook J, Thordal-Christensen H, Katagiri F (2011) Physical association of Arabidopsis hypersensitive induced reaction proteins (HIRs) with the immune receptor RPS2. J Biol Chem 286: 31297–31307

Qiao Z, Kong Q, Tee WT, Lim ARQ, Teo MX, Olieric V, Low PM, Yang Y, Qian G, Ma W, et al (2022) Molecular basis of the key regulator WRINKLED1 in plant oil biosynthesis. Sci Adv 8: eabq1211

Rappsilber J, Mann M, Ishihama Y (2007) Protocol for micro-purification, enrichment, pre-fractionation and storage of peptides for proteomics using StageTips. Nat Protoc 2: 1896– 1906

Rekhter D, Lüdke D, Ding Y, Feussner K, Zienkiewicz K, Lipka V, Wiermer M, Zhang Y, Feussner I (2019) Isochorismate-derived biosynthesis of the plant stress hormone salicylic acid. Science 365: 498–502

Rosenzweig R, Nillegoda NB, Mayer MP, Bukau B (2019) The Hsp70 chaperone network. Nat Rev Mol Cell Biol 20: 665–680

Salvucci ME (2007) Association of Rubisco activase with chaperonin-60: a possible mechanism for protecting photosynthesis during heat stress. J Exp Bot 59: 1923–1933

Schieferle S, Tappe B, Korte P, Mueller MJ, Berger S (2021) Pathogens and elicitors induce local and systemic changes in triacylglycerol metabolism in roots and in leaves of Arabidopsis thaliana. Biology (Basel) 10: 920

Schmid M, Davison TS, Henz SR, Pape UJ, Demar M, Vingron M, Schölkopf B, Weigel D, Lohmann JU (2005) A gene expression map of Arabidopsis thaliana development. Nat Genet 37: 501–506

Schwanhäusser B, Busse D, Li N, Dittmar G, Schuchhardt J, Wolf J, Chen W, Selbach M (2011) Global quantification of mammalian gene expression control. Nature 473: 337–342

Sham A, Moustafa K, Al-Ameri S, Al-Azzawi A, Iratni R, AbuQamar S (2015) Identification of Arabidopsis candidate genes in response to biotic and abiotic stresses using comparative microarrays. PLoS One 10: e0125666

Shevchenko A, Tomas H, Havlis J, Olsen JV, Mann M (2006) In-gel digestion for mass spectrometric characterization of proteins and proteomes. Nat Protoc 1: 2856–2860

Shimada TL, Takano Y, Shimada T, Fujiwara M, Fukao Y, Mori M, Okazaki Y, Saito K, Sasaki R, Aoki K, et al (2014a) Leaf oil body functions as a subcellular factory for the production of a phytoalexin in Arabidopsis. Plant Physiol 164: 105–118

Shimada TL, Takano Y, Shimada T, Fujiwara M, Fukao Y, Mori M, Okazaki Y, Saito K, Sasaki R, Aoki K, et al (2014b) Leaf oil body functions as a subcellular factory for the production of a phytoalexin in Arabidopsis. Plant Physiol 164: 105–118

Shiva S, Samarakoon T, Lowe KA, Roach C, Vu HS, Colter M, Porras H, Hwang C, Roth MR, Tamura P, et al (2020) Leaf lipid alterations in response to heat stress of Arabidopsis thaliana. Plants (Basel) 9: 845

Smith MA, Jonsson L, Stymne S, Stobart K (1992) Evidence for cytochrome *b* 5 as an electron donor in ricinoleic acid biosynthesis in microsomal preparations from developing castor bean (*Ricinus communis* L.). Biochem J 287: 141–144

Sparkes IA, Runions J, Kearns A, Hawes C (2006) Rapid, transient expression of fluorescent fusion proteins in tobacco plants and generation of stably transformed plants. Nat Protoc 1: 2019– 2025

Spoel SH, Koornneef A, Claessens SMC, Korzelius JP, Van Pelt JA, Mueller MJ, Buchala AJ, Métraux J-P, Brown R, Kazan K, et al (2003) NPR1 modulates cross-talk between salicylate- and jasmonate-dependent defense pathways through a novel function in the cytosol. Plant Cell 15: 760–770

Su T, Xu J, Li Y, Lei L, Zhao L, Yang H, Feng J, Liu G, Ren D (2011) Glutathione-indole-3-acetonitrile is required for camalexin biosynthesis in *Arabidopsis thaliana*. Plant Cell 23: 364– 380

Sun Q, Zybailov B, Majeran W, Friso G, Olinares PDB, van Wijk KJ (2009) PPDB, the plant proteomics database at Cornell. Nucleic Acids Res 37: D969–D974

Szalainé Ágoston B, Kovács D, Tompa P, Perczel A (2011) Full backbone assignment and dynamics of the intrinsically disordered dehydrin ERD14. Biomol NMR Assign 5: 189–193

Szklarczyk D, Gable AL, Nastou KC, Lyon D, Kirsch R, Pyysalo S, Doncheva NT, Legeay M, Fang T, Bork P, et al (2021) The STRING database in 2021: customizable protein-protein networks, and functional characterization of user-uploaded gene/measurement sets. Nucleic Acids Res 49: D605–D612

Tarazona P, Feussner K, Feussner I (2015) An enhanced plant lipidomics method based on multiplexed liquid chromatography-mass spectrometry reveals additional insights into cold- and drought-induced membrane remodeling. Plant J 84: 621–633

Taurino M, Costantini S, De Domenico S, Stefanelli F, Ruano G, Delgadillo MO, Sánchez-Serrano JJ, Sanmartín M, Santino A, Rojo E (2018) SEIPIN proteins mediate lipid droplet biogenesis to promote pollen transmission and reduce seed dormancy. Plant Physiol 176: 1531–1546

Tyanova S, Temu T, Sinitcyn P, Carlson A, Hein MY, Geiger T, Mann M, Cox J (2016) The Perseus computational platform for comprehensive analysis of (prote)omics data. Nat Methods 13: 731–740

Tzen J, Cao Y, Laurent P, Ratnayake C, Huang A (1993) Lipids, proteins, and structure of seed oil bodies from diverse species. Plant Physiol 101: 267–276

Uknes S, Winter AM, Delaney T, Vernooij B, Morse A, Friedrich L, Nye G, Potter S, Ward E, Ryals J (1993) Biological induction of systemic acquired resistance in Arabidopsis. Mol Plant Microbe Interact 6: 692–698

Vance VB, Huang AH (1987) The major protein from lipid bodies of maize. Characterization and structure based on cDNA cloning. J Biol Chem 262: 11275–11279

Vizcaíno JA, Deutsch EW, Wang R, Csordas A, Reisinger F, Ríos D, Dianes JA, Sun Z, Farrah T, Bandeira N, et al (2014) ProteomeXchange provides globally coordinated proteomics data submission and dissemination. Nat Biotechnol 32: 223–226

Wang K, Senthil-Kumar M, Ryu C-M, Kang L, Mysore KS (2012) Phytosterols play a key role in plant innate immunity against bacterial pathogens by regulating nutrient efflux into the apoplast. Plant Physiol 158: 1789–1802

Wang T-Y, Wu J-R, Duong NKT, Lu C-A, Yeh C-H, Wu S-J (2021) HSP70-4 and farnesylated AtJ3 constitute a specific HSP70/HSP40-based chaperone machinery essential for prolonged heat stress tolerance in Arabidopsis. J Plant Physiol 261: 153430

Welti R, Li W, Li M, Sang Y, Biesiada H, Zhou H-E, Rajashekar CB, Williams TD, Wang X (2002) Profiling membrane lipids in plant stress responses. Role of phospholipase D alpha in freezing-induced lipid changes in Arabidopsis. J Biol Chem 277: 31994–32002

Wilkins O, Bräutigam K, Campbell MM (2010) Time of day shapes Arabidopsis drought transcriptomes. Plant J 63: 715–727

Xu C, Fan J, Froehlich JE, Awai K, Benning C (2005) Mutation of the TGD1 chloroplast envelope protein affects phosphatidate metabolism in *Arabidopsis*. Plant Cell 17: 3094–3110

Xu C, Fan J, Riekhof W, Froehlich JE, Benning C (2003) A permease-like protein involved in ER to thylakoid lipid transfer in Arabidopsis. EMBO J 22: 2370–2379

Xu C, Shanklin J (2016) Triacylglycerol metabolism, function, and accumulation in plant vegetative tissues. Annu Rev Plant Biol 67: 179–206

Yamaguchi Y, Nakamura T, Kusano T, Sano H (2000) Three Arabidopsis genes encoding proteins with differential activities for cysteine synthase and β-cyanoalanine synthase. Plant Cell Physiol 41: 465–476

Yao J, Withers J, He SY (2013) Pseudomonas syringae infection assays in Arabidopsis. Methods Mol Biol 1011: 63–81

Yu L, Zhou C, Fan J, Shanklin J, Xu C (2021) Mechanisms and functions of membrane lipid remodeling in plants. Plant J 107: 37–53

Zhu J-K (2016) Abiotic stress signaling and responses in plants. Cell 167: 313–324

